# Structural basis of *Fusobacterium nucleatum* adhesin Fap2 interaction with receptors on cancer and immune cells

**DOI:** 10.1101/2024.02.28.582045

**Authors:** Felix Schöpf, Gian L. Marongiu, Klaudia Milaj, Thiemo Sprink, Judith Kikhney, Annette Moter, Daniel Roderer

## Abstract

The intestinal microbiome (IM) is decisive for the human host’s health. Numerous microbiota drive the progression of colorectal cancer (CRC), the third-most common cancer worldwide. The Gram-negative *Fusobacterium nucleatum* (Fn) is overrepresented in the IM of CRC patients and has been correlated with the emergence, progression, and metastasis of tumors. A key pathogenic factor of Fn is the adhesin Fap2, an autotransporter protein that facilitates association to cancer and immune cells via two receptors, the glycan Gal-GalNAc and the T-cell protein TIGIT, respectively. The latter interaction leads to deactivation of immune cells. Mechanistic details of the Fap2/TIGIT interaction remain elusive due to the lack of high-resolution structural data. Here, we report a system to recombinantly express functional Fap2 on the *Escherichia coli* surface, which interacts with Gal-GalNAc on cancer cells and with purified TIGIT with submicromolar affinity. Cryo-EM structures of Fap2, alone and in complex with TIGIT, show that the ∼50 nm long rod-shaped Fap2 extracellular region binds to TIGIT on its membrane-distal tip via an extension of a β-helix domain. Moreover, by combining structure predictions, cryo-EM, docking and MD simulations, we identified a binding pit for Gal-GalNAc on the tip of Fap2. Our data represent the first purification and high-resolution structural analysis of a Fn autotransporter adhesin and its receptor association.

## Introduction

The human intestinal microbiome (IM) comprises up of 10^14^ microbes, a higher number than human cells in the body (Sender, Fuchs and Milo, 2016). With estimated 22 million genes (Tierney et al., 2019), the genetic diversity of the IM is orders of magnitude higher than the human genome. The composition of the IM contributes largely to the host’s metabolism and health by providing different secreted bacterial enzymes, metabolites and lipids (Wu et al., 2021; Lamichhane et al., 2021). Pathological modulations of the IM, termed dysbiosis, have a severe effect on the intestinal system’s health. Such dysbiosis is one of the key determinants for the prevalence of colorectal cancer (CRC), the third most common cancer worldwide and the second most cause of cancer related death (Montalban-Arques and Scharl, 2019). 1.9 million new CRC cases and 935,000 deaths were estimated globally for 2020 (Sung et al., 2021), with incidences consistently rising worldwide (Arnold et al., 2017, 2020). The over-representation of particular microbes in the IM can cause the onset and drive progression of CRC through various mechanisms, such as causing DNA damage through toxins (Wilson et al., 2019), growth stimulation of tumor cells through the activation of signal transduction pathways, or immune system down-regulation (Levy et al., 2017). *Fusobacterium nucleatum* (Fn), an anaerobic Gram-negative rod, has been demonstrated to act via the latter two mechanisms (Rubinstein et al., 2013; Kostic et al., 2013; Rubinstein et al., 2019). Fn has been found over-represented in the IMs of CRC patients in numerous studies (Castellarin et al., 2012; Kostic et al., 2012; Ito et al., 2015; Dai et al., 2018; Wirbel et al., 2019; Ai et al., 2019; Loftus, Hassouneh and Yooseph, 2021), and Fn has therefore been suggested as a biomarker for CRC (Yuan et al., 2021). Moreover, Fn has been shown to drive breast cancer growth (Parhi et al., 2020), and to be associated with preterm birth, stillbirth (Han et al., 2004), and periodontal disease (Liu et al., 2010), highlighting the high clinical relevance of this understudied pathogen. In contrast to other important CRC driving microbiota, e.g., genotoxic *Escherichia coli* and *Bacteroides fragilis* (Valguarnera and Wardenburg, 2020), Fn encodes no known secreted toxins (Brennan and Garrett, 2019). Instead, Fn encodes numerous outer membrane (OM) anchored adhesins that are utilized to attach to other microbes in biofilms (Kaplan et al., 2009; Lima, Shi and Lux, 2017) or directly to host cells (Xu et al., 2007; Coppenhagen-Glazer et al., 2015; Shhadeh et al., 2021). The latter causes tumor growth stimulation, such as by interaction of the adhesin FadA to E-cadherin and subsequent activation of β-catenin translocation to the nucleus (Rubinstein et al., 2013; Guo et al., 2020), or by interaction with the T-cell immunoreceptor with immunoglobulin and ITIM domains (TIGIT) on T cells or natural killer (NK) cells via the adhesin Fap2 (Gur et al., 2015).

Fap2 is a type Va autotransporter OM protein (Kaplan et al., 2005) and with ca. 400 kDa the largest single protein in Fn (Sanders et al., 2018). It is a bifunctional adhesin through which Fn binds to tumors via Gal-GalNAc (Abed et al., 2016), a glycan that is abundant on and specific for cancer cells in the colon (Yang and Shamsuddin, 1996), and to immune cells, specifically NK cells, via TIGIT (Gur et al., 2015). The combination of both interactions make Fap2 a key molecule in Fn-driven CRC, as it mediates tumor colonization independently of the level of E-cadherin expression on tumor cells (Abed et al., 2016) as well as clearance of the tumor from active NK cells (Zhang et al., 2018). The molecular mechanisms of how Fap2 binds to the two receptors remain however unknown because no high-resolution structure of Fap2 or another Fn autotransporter adhesin is known.

We have therefore developed an expression system for the recombinant production of Fap2 from the clinically relevant Fn strain ATCC23726 on the OM surface of *E. coli*, which we successfully applied to produce functional Fap2 that binds to its two receptors, Gal-GalNAc and TIGIT, and thereby facilitates bacterial association to these cells. Structural analysis of the Fap2 extracellular domain (ECD) using cryogenic electron microscopy (cryo-EM) and single particle analysis (SPA) in combination with AlphaFold2 structure prediction reveals a rod-shaped 46,5 nm long β-helix with a matchstick-like tip that interacts with the immune cell receptor TIGIT. Molecular docking and MD simulations indicate that the cancer cell receptor Gal-GalNAc binds to the back side of the matchstick tip. We therefore conclude that tumor-colonizing Fn uses the elongated shape of Fap2 in a spear-like manner to keep NK cells away from the tumor, whereas the adhesin simultaneously acts as a hook to remain attached to tumor cells. Our recombinant expression system can be adjusted to express further fusobacterial adhesins, omitting the necessity for anaerobic work with Fn, and has a general transfer potential to the production of other bacterial surface proteins.

## Results

### Production of the Fap2-ECD in fusion with an *E. coli* autotransporter

The genome of *F. nucleatum subsp. nucleatum* ATCC23726 contains 2111 ORFs (Sanders et al., 2018), 21 of them containing autotransporter domains as annotated by InterProScan5 (Jones et al., 2014). These are on average larger proteins than other Fn ORFs, notably the largest two of them are the arginine-binding protein RadD and the virulence factor Fap2 (Fig. 1a) (Sanders et al., 2018). It had been shown that these two autotransporter adhesins are strongly expressed in Fn and localized in the OM (Kaplan et al., 2010). Despite their importance in biofilm formation and attachment to the human host, no molecular characterization of either adhesin has been reported, probably limited by the lack of access to the purified proteins so far. As an obligate anaerobe and pathogen, cultivation and handling of Fn is restricted to biosafety level 2 laboratories with anaerobic cultivation conditions. This hampers the analysis under our experimental conditions, especially the co-culture with aerobic cell lines. To facilitate the latter, we applied the AIDA autotransporter system to recombinantly produce Fap2 on the OM surface of *E. coli* using pAIDA1 (Lattemann et al., 2000; Jarmander et al., 2012) as template. Domain analysis indicated that the Fap2-ECD contains 3430 out of 3744 residues of the mature protein, followed by a C-terminal, 314 residues long autotransporter domain (Fig. 1b). To identify the regions of the ECD to be attached to the AIDA autotransporter, we predicted the structure of full Fap2 using AlphaFold2 (Jumper et al., 2021). This revealed a 47 nm long β-helix that comprises 2905 residues, where 325 residues close to the N-terminus form a region with low prediction confidence (SI Fig. 1a). The β-helix is followed by a linker of 200 residues, which connects the β-helix to the autotransporter domain (Fig. 1c). While the autotransporter, the linkers, and the β-helix are highly conserved in different Fn strains, more variations are evident at the N-terminal part between Fap2 of Fn ATCC23726 and Fn ATCC25586 (SI Fig. 1b).

**Figure 1:**
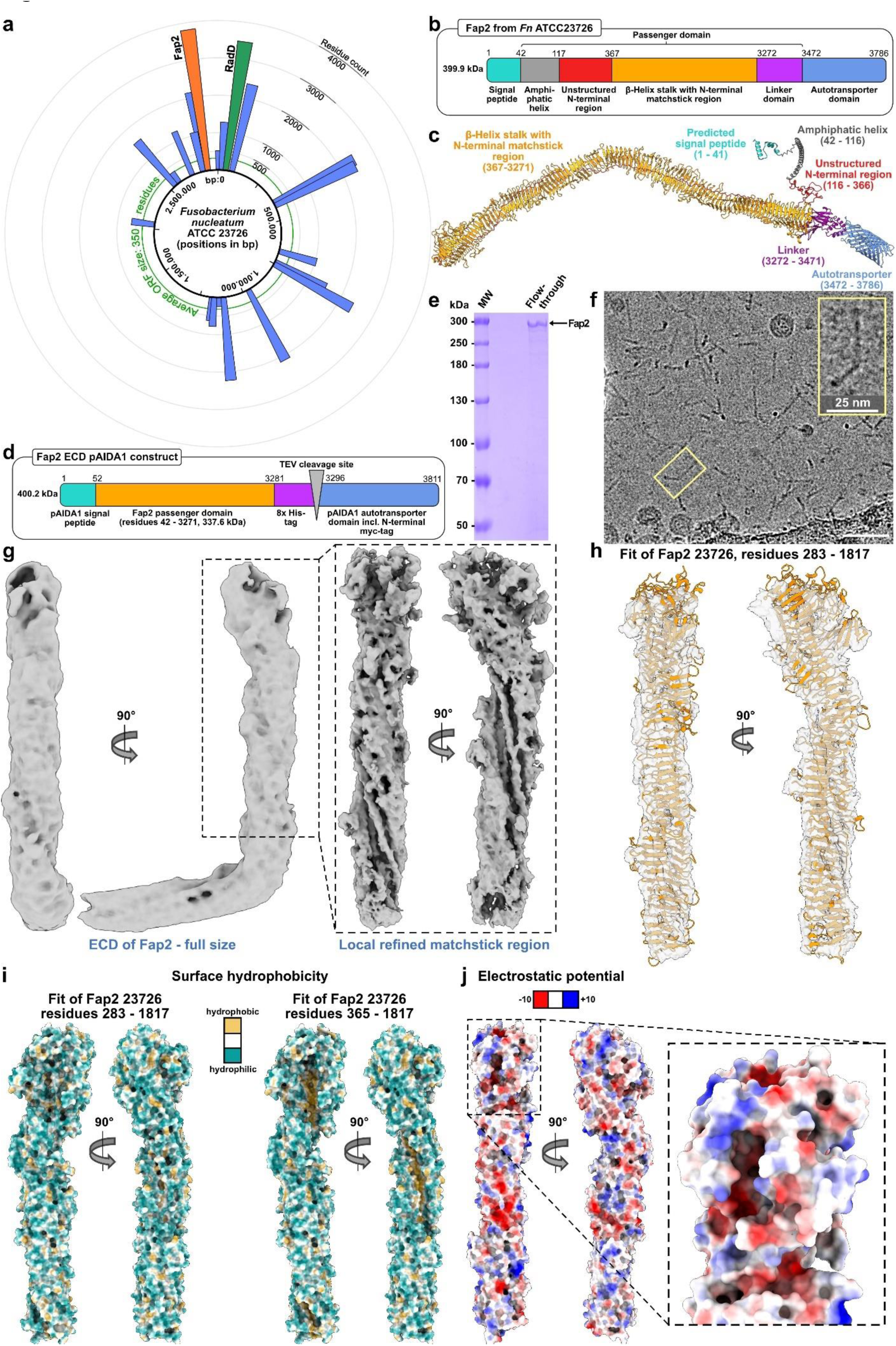
Domain analysis, recombinant production and structure of Fap2. a) Genome organization of Fn ATCC23726 with identified sequences for autotransporter adhesins (blue), in which Fap2 and RadD are indicated in orange and green, respectively. b) Domain organization of Fap2, as identified by AlphaFold2. c) AlphaFold2 structure prediction of Fap2 from Fn ATCC23726, colored by domain as in (b). d) Design of the pAIDA_TEV_Fap2EC expression plasmid, which contains the unstructured N-terminal region, the matchstick and the β-helix of Fap2 (residues 42 - 3271). e) SDS-PAGE, showing purified Fap2-ECD after release from the AIDA autotransporter, Ni-NTA purification and removal of GST-TEV. f) Section of a Cryo-EM micrograph of purified Fap2-ECD, recorded at 300 kV and 2.8 µm defocus using a Falcon 3 detector. Scale bar: 50 nm. g) Cryo-EM density maps of Fap2-ECD at 4.7 Å resolution (left panel), and local refinement of the tip of the longer branch, which displays a hook-like end (right panel). h) Density map with residues 283-1817 of the matchstick region and the β-helix of Fap2 fitted into the longer branch. i) Surface representation of the longer branch of Fap2 colored by hydrophobicity, in which the longitudinal groove is shown with (left) and without (right) fitted unstructured N-terminus. j) Surface representation of the longer branch of Fap2 colored by electrostatics at pH 7. The box highlights the matchstick region with a hollow pit.

Since pAIDA1 contains a signal sequence and also a linker that are both designed to enable translocation of heterologous cargos through the AIDA autotransporter, we cloned the variable N-terminal part and the entire β-helix without the Fap2 linkers between the pAIDA signal sequence and the linker. To facilitate the isolation of pure Fap2-ECD, we introduced a His-tag before the tobacco mosaic virus (TEV) protease cleavage site between the Fap2-ECD and AIDA autotransporter (pAIDA_TEV_Fap2EC, Fig.1d). We then expressed pAIDA_TEV_Fap2EC recombinantly in *E. coli* and indeed observed the presence of our fusion protein in the membrane fraction (SI Fig. 2), indicating successful translocation of the Fap2-ECD through the AIDA autotransporter domain. After TEV cleavage, Ni^2+^ affinity chromatography, and removal of TEV protease by glutathione affinity chromatography, we obtained the 340 kDa large Fap2-ECD at >90% purity (Fig. 1e, SI Fig. 2b-g), suitable for structural and biophysical analysis. In agreement with the β-helix of the AlphaFold2 prediction, the protein forms up to about 50 nm long rods, as evident in cryo-EM micrographs (Fig. 1f).

**Figure 2:**
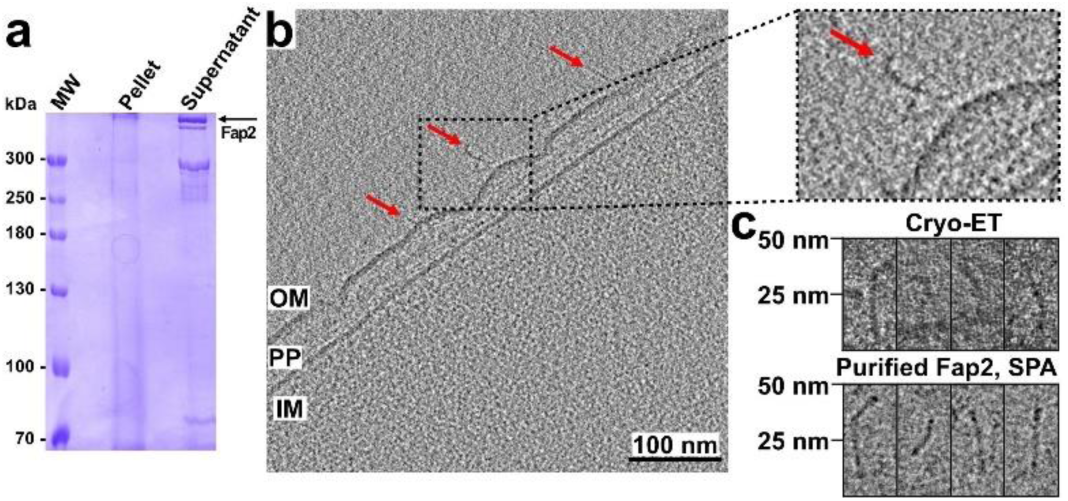
Comparison of purified Fap2-ECD with native Fn outer membrane adhesins. a) SDS-PAGE of the isolated OM fraction of Fn ATCC25586 after co-cultivation with Jurkat cells. Proteins corresponding to the indicated bands were identified as Fap2 and RadD in peptide fingerprint analysis (SI Figure 5a,b). b) Representative cryo tomogram of a section of Fn ATCC25586 including the inner membrane (IM), periplasm (PP) and outer membrane (OM). Rod-like structures on the surface that represent autotransporter adhesins in size and shape are indicated by arrowheads. The inset shows one of these magnified. c) Selected rod-shaped OM surface protrusions from Fn (upper row) in comparison with different projection views of the recombinantly produced Fap2-ECD (lower row).

Cryo-EM and SPA of the Fap2-ECD at an overall resolution of 4.7 Å (SI Fig. 3, SI Table 1) revealed a total length of 45 nm and width of 4,5 nm for the ECD. The individual strands of the β-helix are not entirely separated in the present density map, however a triangular-shaped groove of 2 nm width along the longitudinal axis is evident (Fig. 1g, SI Movie 1). The β-helix domain is parted in two branches of 26 and 19 nm, respectively, by a kink of 101°. The longer, membrane-distal branch has a thickening by 1,2 nm, which we term matchstick region. To improve the low local resolution in this region, we carried out signal subtraction of the shorter branch and local refinement of the tip of the longer branch of the ECD (SI Fig. 3,4). With this, we identified a hook-like structure with two triangular extensions (Fig. 1g). These shapes fit the lateral extensions at the membrane-distal tip of the β-helix in the AlphaFold model (residues K525 - K836), and thus allowed fitting of the predicted β-helical structure into the cryo-EM density (Fig. 1h). The map and the model reveal a unique hydrophobic groove that winds around the longitudinal axis of Fap2. This gap is closed by the unstructured proline-rich, and amphiphilic N-terminus (residues N195 - S361), which together result in a hydrophilic surface (Fig. 1h,i, SI Movie 1). The unstructured N-terminus is highly conserved in different fusobacterial species, highlighting also the conservation of the hydrophobic groove, and in predicted autotransporter proteins of the *Pseudoleptotrichia goodfellowii* and *Leptotrichia* species that are known human pathogens (SI Fig. 1a, 3i). No significant sequence similarity was identified outside the order *Fusobacterales*, indicating that this particular structural motif is so far specific for the adhesins of these bacteria. The triangular extensions at the matchstick region result in two pits at the tip (Fig. 1j), which are present only in Fap2, but not in RadD (SI Fig. 1a) and are probably involved in receptor binding.

**Figure 3:**
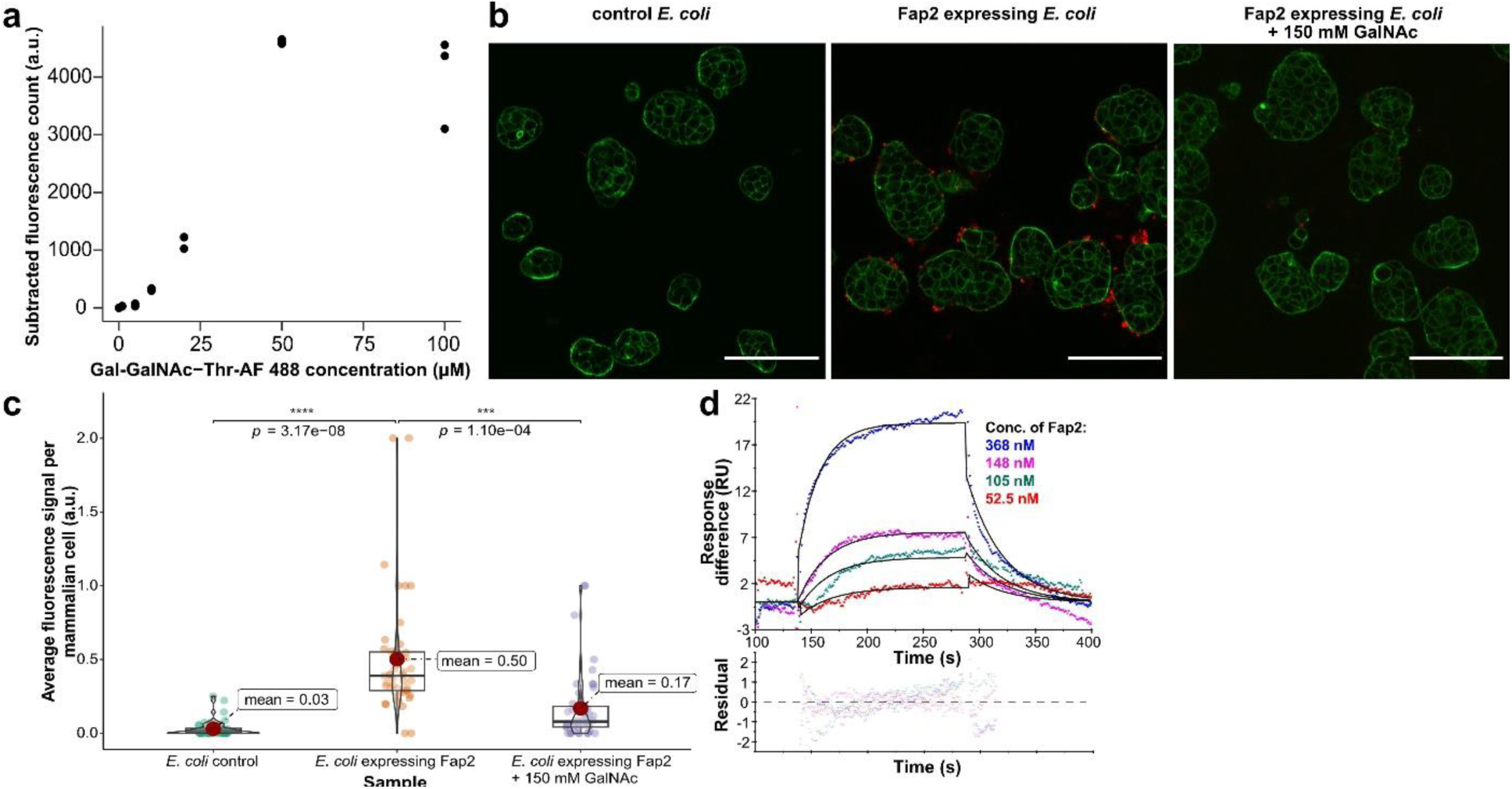
Adhesion of Fap2 to Gal-GalNAc-Thr and hTIGIT. a) Concentration-dependent binding of fluorescently labeled Gal-GalNAc-Thr to Fap2-expressing *E. coli* and control *E. coli* that express pAIDA without the Fap2-ECD. Data points represent three replicates per concentration. b) Fluorescence micrographs that show cancer cell binding (HT-29) of Fap2-expressing *E. coli*. Cell binding is reduced in the presence of an excess of GalNAc (right panel). Red: *E. coli* labeled with CellBrite Fix 555, green: actin labeled with Phalloidin-iFluor 488. Scale bar: 100 µm. Note the unequal distribution of *E. coli* on HT-29 cells. c) Quantification of experiments as in b), with average fluorescence signal of *E. coli* bound to HT-29 determined from 50 micrographs each. The mean is shown as red dot. The boxes represent the interquartile range, with the median shown as horizontal line within the box. In addition, the distribution of data points is visually represented by violin plots. d) Quantification of Fap2-ECD binding to hTIGIT-ECD using surface plasmon resonance. A global fit according to a 1:1 binding model (black solid lines) reveals a K_D_ of ∼580 nM (SI Table 2). Lower panel: residuals of fit.

**Figure 4:**
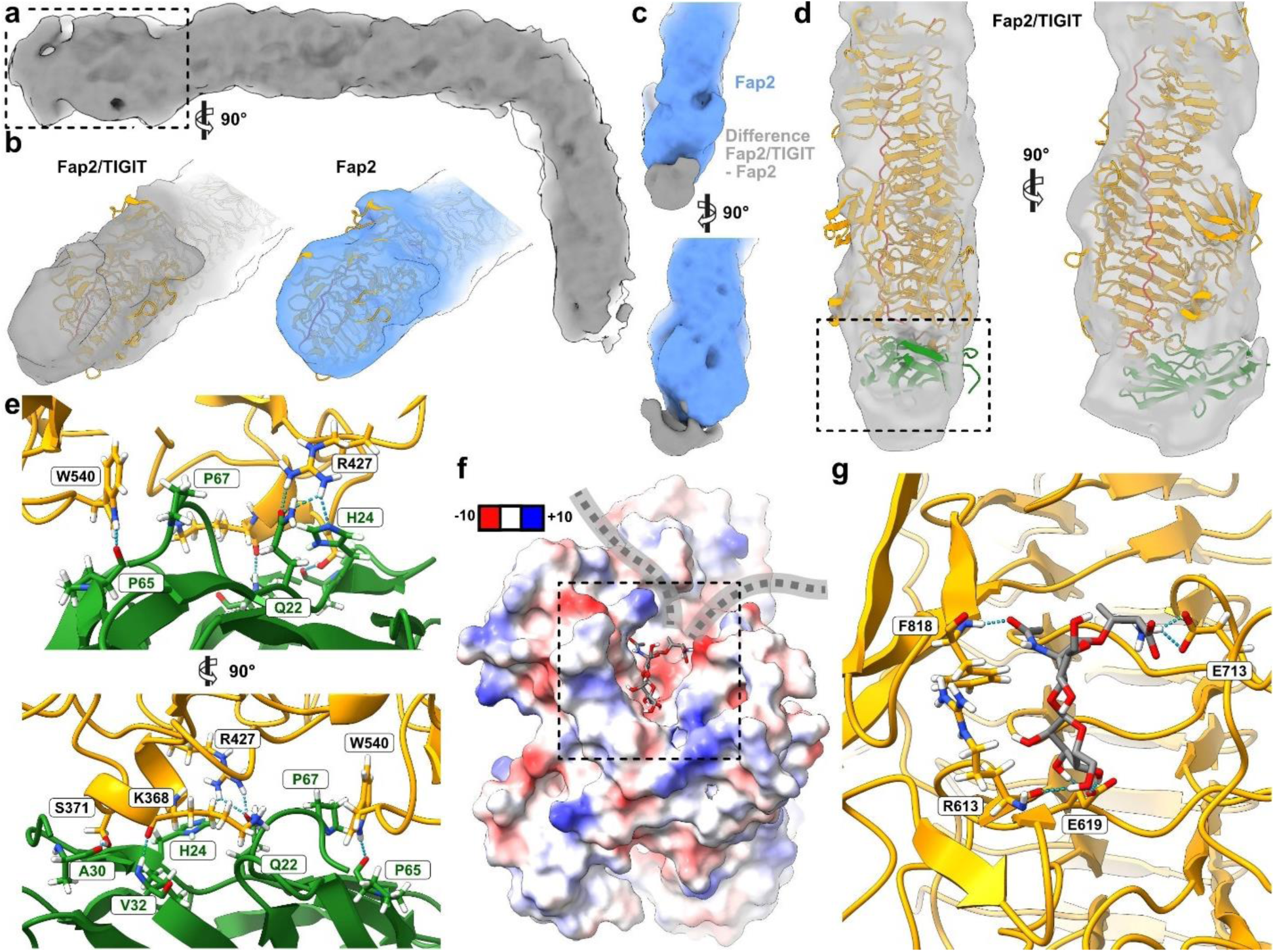
Structure of the complex of Fap2-ECD and hTIGIT-ECD and docking of Gal-GalNAc-Thr. a) Cryo-EM density map of Fap2-ECD/hTIGIT-ECD at 6.0 Å resolution. b) Comparison of the cryo-EM densities of the membrane-distal tip regions of Fap2-ECD/hTIGIT-ECD (left) and Fap2-ECD (right), as indicated by the box in a). The Fap2 model is shown in orange for both maps. c) Difference density of Fap2-ECD/hTIGIT-ECD and Fap2-ECD (gray) shown atop the tip of Fap2-ECD (blue). Difference density was obtained through subtraction of lowpass-filtered maps (10 Å) and is shown after further lowpass filtering to 12 Å. d) Docking of hTIGIT-ECD (green) to Fap2-ECD (orange) using HADDOCK (van Zundert et al., 2016). The best docking pose is shown and is in agreement with the cryo-EM density. e) Proposed interaction site of Fap2 and hTIGIT as indicated by the box in d). The loop G66-P67-G68 of hTIGIT intercalates into a gap in Fap2, and main chain and side chain interactions as identified in docking and verified by MD simulations are highlighted. f) Surface representation of Fap2 colored according to Coulomb potential (red: negative, blue: positive charges) with docked Gal-GalNAc-Thr. A representative frame of the MD simulation of the best docking pose is shown (SI Movie 4). The Fap2 model has two clefts adjacent to the docking sites, which is in agreement with the polypeptide chain of the Gal-GalNAc-containing O-glycosylated receptor going through (indicated by gray dashed line). g) Hydrogen bonds between the docked Gal-GalNAc-Thr and R613, E619, E713, and F818 of Fap2, which remain most consistent in MD simulation.

Our findings report the first high-resolution structural analysis of Fap2 as a fusobacterial autotransporter adhesin and its receptor-binding β-helical ECD. We conclude that through the long rod-like extracellular domain Fap2 acts in functional analogy to similarly shaped autotransporter adhesins, such as CdrA/B from *Pseudomonas aeruginosa* (Melia et al., 2021).

### Shape and size of recombinantly produced Fap2 resemble adhesins in the OM of Fn

To compare recombinantly purified Fap2 with native autotransporter adhesins in the Fn OM, we cultivated Fn ATCC25586 in the presence of the immortalized T-cell line Jurkat, which had been demonstrated to increase Fap2 production in Fn ATCC23726 (Kaplan et al., 2010). We indeed observed a strong expression of several > 300 kDa proteins in the OM fraction that make up more than 50% of all solubilized Fn OM proteins, as evident in SDS-PAGE (Fig. 2a). Peptide fingerprint analysis of the bands indicated that the three upper bands represent Fap2, as reported for Fn ATCC23726 previously (Kaplan et al., 2010), while the lower band predominantly contains RadD (SI Fig. 5a,b), both of them being the largest ORFs in Fn (Fig. 1a). To analyze the adhesins on the native OM surface, we vitrified Fn ATCC25586 and visualized their surfaces using cryo electron tomography (cryo-ET). In fact, we could identify kinked rod-like structures protruding from the OM (Fig. 2b). These cell attachments appear up to 50 nm long, which matches the lengths of recombinantly produced Fap2-ECD as evident in cryo-EM micrographs (Fig. 2c). Moreover, the membrane-distal ends of some cell attachments display bulges, as observed in the cryo-EM density of the longer branch of Fap2-ECD (Fig. 1g, Fig. 2b,c). After detergent extraction and isolation from the OM, the cell attachments keep their shape and integrity, as indicated by SDS-PAGE and negative stain EM (SI Fig. 5c-e).

**Figure 5:**
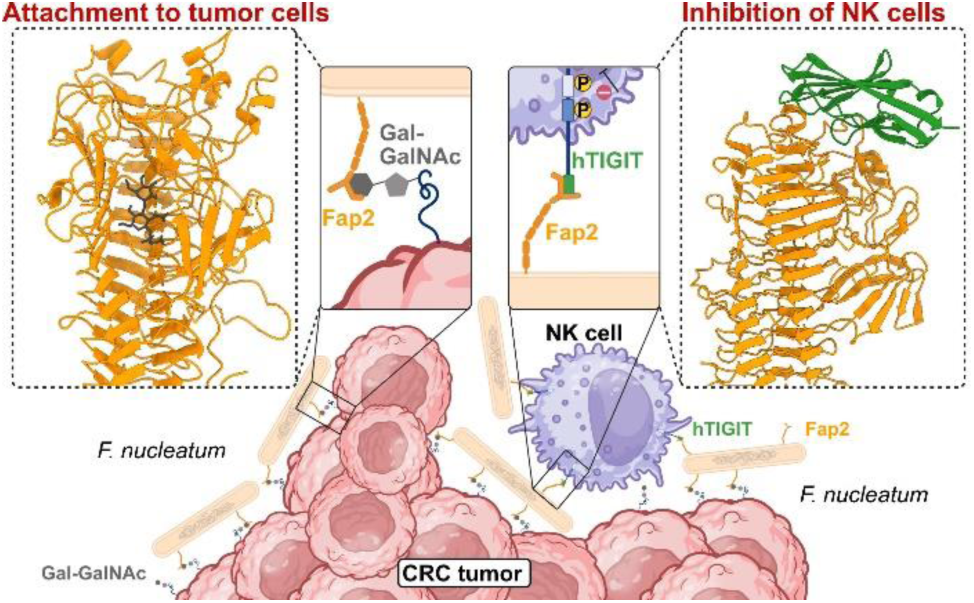
Proposed mechanism of how Fn applies Fap2 to associate to tumors and deactivate tumor-invading NK cells through interaction with Gal-GalNAc and hTIGIT on the back side and front side of the Fap2 matchstick tip, respectively.

Taken together, our findings suggest that a large number of Fap2 and RadD molecules are present in the Fn OM, where the linker domains allow flexible attachment. Therefore, the entire flanks and tip of the rod-like β-helix would be accessible for interaction with Gal-GalNAc and TIGIT by Fap2.

### Recombinantly produced Fap2 facilitates tumor cell attachment via Gal-GalNAc

Having verified successful recombinant production of Fap2 that resulted in protein with native-like shape and domain arrangement, we set out to test the functionality of the recombinant Fap2-ECD, i.e., the association with its two known receptors, Gal-GalNAc and hTIGIT. The disaccharide Gal-GalNAc is an abundant O-glycosylation of surface proteins of colon cancer cells, thereby serving as biomarker for CRC (Yang and Shamsuddin, 1996), and facilitates tumor colonization of Fn in orthotopic mouse models (Abed et al., 2016). To examine whether recombinant Fap2 on the *E. coli* surface binds to Gal-GalNAc, we transformed the bacteria with pAIDA_TEV_Fap2EC, induced expression, and incubated with a fluorescently labeled Gal-GalNAc-Thr adduct (SI Fig. 6a-c). In fact, we observed increased Gal-GalNAc binding for Fap2-expressing *E. coli* compared to *E. coli* expressing pAIDA without Fap2 at receptor concentrations as low as 25 - 100 µM (Fig. 3a), exceeding previously reported concentrations in the mM range (Abed et al., 2016). This means that recombinant Fap2 binds to the cancer cell receptor at least as well as native Fap2. Next, we addressed whether recombinant Fap2 facilitates cancer cell adhesion of *E. coli*. For this, we stained the membranes of *E. coli*, and analyzed its association to the CRC cell line HT-29 via fluorescence microscopy. Indeed, we identified clusters of Fap2-expressing *E. coli* attached to the surface of HT-29, which are less abundant in Fap2-negative bacteria (Fig. 3b,c, SI Fig. 6d). Notably, the bacteria are not evenly distributed among cells, but cluster at higher numbers on some, whereas other cells are practically unoccupied. This indicates that surface Gal-GalNAc is not equally accessible to the bacteria within the cell population. Similar observations were made for Caco-2 CRC cells (SI Fig. 6e). Our experiments demonstrate that recombinant Fap2 has sufficient functional affinity to allow association with cancer cells, even those with low Gal-GalNAc surface levels such as HT-29 (Abed et al., 2016), and that no other *E. coli* surface proteins bind to a surface receptor on the CRC cells with an affinity comparable to that of Fap2 for Gal-GalNAc. To verify that Fap2-expressing *E. coli* attach to Gal-GalNAc and not to further possible receptors on cancer cells, we carried out a competition assay with GalNAc in solution. As expected, attachment to HT-29 was tremendously reduced in the presence of 150 mM soluble GalNAc almost to the level of non-induced bacteria (Fig. 3b,c). We therefore conclude that the interaction is specific between Gal-GalNAc and Fap2, involving no other receptors of *E. coli* and HT-29 cells. Together, our results demonstrate that Fap2 attached to the *E. coli* autotransporter AIDA is functional and allows binding to CRC cells. Therefore, we present a simplified *in cellulo* system that allows to assess Fn cell adhesion in its natural context without the need for anaerobic work or higher biosafety levels, enabling co-culture experiments with oxygen-dependent human cell lines and clinically relevant experiments to a broader research community.

### Recombinantly produced Fap2 interacts with TIGIT

After confirming functional binding of recombinant Fap2 to cancer cells via the glycan receptor Gal-GalNAc, we next addressed its association to the receptor on immune cells, TIGIT (Gur et al., 2015). To study the interaction of Fap2 with native-like, glycosylated human TIGIT (hTIGIT), we expressed the extracellular domain of human hTIGIT (residues 1 – 141; hTIGIT-ECD) together with a C-terminal eGFP and StrepII tag in Expi cells and purified the protein (SI Fig. 7a-c). The blurred band of purified hTIGIT-ECD on SDS-PAGE indicates that the protein is glycosylated at both predicted sites (N32 and N101) (Lin et al., 2021), and we proceeded to test the interaction of purified hTIGIT-ECD with Fap2.

We then carried out a qualitative pulldown assay for which we immobilized Fap2-ECD on NiNTA and loaded secreted hTIGIT as prey. Indeed, we identified hTIGIT-ECD as binding partner (SI Fig. 7d), whereas non-glycosylated hTIGIT purified from *E. coli* (Stengel et al., 2012) did not bind to Fap2 (SI Fig. 7e,f). This indicates the direct or indirect involvement of one or both glycans in the binding reaction, similar as reported for the interaction of hTIGIT with the poliovirus receptor (PVR) (Lin et al., 2021). To evaluate the physiological relevance of the Fap2-hTIGIT interaction in comparison to other known hTIGIT interaction partners (PVR and CD112), we set out to quantify the affinity of glycosylated hTGIT-ECD and Fap2-ECD using surface plasmon resonance (SPR). The obtained dissociation constant (K_D_) of ∼580 nM (Chi^2^ = 0.477, k_on_ = 52,900 ± 3,120 M^-1^ s^1^, k_off_ = 0.03 ± 0,001 s^-1^; Fig. 3d, SI Table 2) is between those reported for hTIGIT and PVR of 1-3 nM (Yu et al., 2009; Saha et al., 2023) and for hTIGIT and Nectin-2 of 5 µM (Deuss et al., 2017). Given the fact that NK cells are deactivated via hTIGIT interaction to PVR (Stanietsky et al., 2009), but also via interaction with the low-affinity receptor Nectin-2, both of them strongly expressed on the surface of cancer cells (Lozano et al., 2020), we conclude that the Fap2 affinity to hTIGIT is suffcient for NK cell deactivation independent of cancer cell receptor binding of hTIGIT.

### hTIGIT binds to the membrane-distal tip of Fap2

To identify where and how Fap2 binds to hTIGIT, we vitrified the complex of Fap2-ECD and hTIGIT-ECD after pulldown (SI Fig. 7d, SI Fig. 8a) for cryo-EM and SPA. The density map with a global resolution of 6.0 Å (SI Fig. 8) revealed a long, kinked, rod-like structure, similar to the Fap2-ECD alone (Fig. 4a). Notably, the tip of the longer branch of the rod has an additional, angular-shaped density, whereas the tip of Fap2 alone appears round-shaped. The density at the tip of the map appeared fragmented because of a mixture of Fap2 with or without hTIGIT. We therefore proceeded with focused 3D classification on the tip and combined subsets with clearly defined density beyond the Fap2 tip, resulting in 45,207 out of 73,482 particles for further refinement (Fig. 4b, SI Fig. 8e,f). After density map subtraction of Fap2 alone, a cuboid shaped density of 5 nm length and 3 nm width at the topmost part of the Fap2 β-helix remained, which is large enough to accommodate one hTIGIT-ECD (Fig. 4c).

Due to the limited resolution of the cryo-EM map, we applied an integrative approach to model the hTIGIT-ECD (residues 23 - 128) on Fap2. For this, we first identified residues of Fap2 that are in close spatial proximity to the additional density in the complex and then docked the polypeptide part of hTIGIT-ECD (PDB 3UCR) to this Fap2 part using HADDOCK (van Zundert et al., 2016). The best cluster of docking poses (score: -14.6 +/-6.7) revealed that loop P87-G92 on hTIGIT-ECD intercalates into a cleft at the Fap2 tip that is formed between D360, T397, R398, E425, N494, D496, and W540 (Fig. 4d,e). We then validated the interaction in molecular dynamics (MD) simulations and confirmed a stable interaction of the complex as suggested by docking, with hTIGIT remaining bound throughout the trajectory in all 10 replicates (SI Movie 2,3). Fap2-W540 forms a hydrogen bond with hTIGIT-P65 and simultaneously interacts via hydrophobic contacts with P67, which intercalates deeply into the cleft at the Fap2 tip. Hydrogen bonds between Fap2-W540, R427, S371 and K368, and hTIGIT-Q22, H24 A30, V32 and P65 are identified as the major contributors to the interaction (Fig. 4e, SI Table 4). Fap2-R427 forms a consistent hydrogen bond with the hTIGIT-Q22 and H24 side chains (SI Movie 2). Fap2-N541, S371 and K368 form a C-shaped interaction interface and participate in consistent hydrogen bonds with hTIGIT-W53, A30 and V32, respectively, throughout all MD simulation replicates (SI Movie 3).

The Fap2/hTIGIT interface is on the same side as the hTIGIT dimerization interface, as evident in the co-structure of hTIGIT/PVR (PBD 3UDW), but differs considerably with respect to orientation and surface accessibility (SI Fig. 8g). Both glycosites (N32 and N101) protrude away from Fap2 in the complex, however, N32 is in close spatial proximity to a predominantly hydrophilic gap between Fap2 and hTIGIT (SI Fig. 8h). This suggests an indirect role for the hTIGIT glycans in interacting with Fap2, probably by introducing slight structural changes in the intercalating loop.

To get insight on the binding site of the CRC cell receptor Gal-GalNAc and whether both receptors can bind to Fap2 simultaneously, we docked the Gal-GalNAc disaccharide attached to Thr, as present in many O-glycosylations, to Fap2-ECD. Half of the docking poses, including the best one (score of -6.1 kcal/mol), were placed in a pit that is located at the back side of the Fap2 matchstick region, opposite to the hTIGIT binding site, facing away from the tip (Fig. 4f, SI Fig. 9a). Notably, this pit shows two openings that would enable the entry and exit of the polypeptide part of O-glycosylated receptors. The Thr attached to Gal-GalNAc in the obtained docking pose aligns with the openings (Fig. 4f), which would be consistent with Fap2 hanging on a Gal-GalNAc-glycosylated polypeptide like a hook on a chain. To identify whether more Gal-GalNAc binding sites exist on the Fap2 β-helix, we carried out further dockings of a larger area of the Fap2-ECD, including the entire matchstick region and the largest part of the longer β-helix branch. Moreover, we docked Gal-GalNAc-Thr to the analogous region of the predicted RadD model (SI Fig. 1a). In Fap2, the highest scores consistently resulted from poses that were docked into the same pit on the matchstick in a similar orientation (SI Fig. 9b). In contrast, docking to RadD resulted in a high variance of different poses (SI Fig. 9c). This shows that the Gal-GalNAc binding pit is specific for Fap2, and is consistent with the absence of such a pit in the predicted model of the otherwise structurally very similar RadD (SI Fig. 9d). To confirm the relevance of docking and analyze the role of individual H-bond interactions, we carried out MD simulations of the docked Gal-GalNAc-Thr-Fap2 complex.

Gal-GalNAc-Thr remained bound and within the pit in 6 out of 10 replicates (SI Movie 4), adopting slightly varying conformations. Hydrogen bond analysis between Fap2 and Gal-GalNAc-Thr of trajectories where the ligand remained bound revealed Fap2-F818, E619 and E713 as major contributors to ligand binding (SI Movie 4). E713 undergoes H-bonds with the Thr amino group of Gal-GalNAc-Thr and not with the glycan moiety, therefore this interaction might differ largely in a Gal-GalNAc-containing polypeptide. The most stable conformation observed showed GalNAc O5 interacting with the F818 backbone amino group. The backbone carbonyl group of R613 interacts with Gal O8; and E619, located deep inside the binding pit, consistently forms hydrogen bonds with Gal O5/O6, which appears to be a prerequisite for stable glycan binding (Fig. 4g, SI Table 5).

The hTIGIT interaction site of Fap2 is ∼3 nm away from the Gal-GalNAc binding pit, which indicates that both receptors have their unique and independent binding sites. Additionally, we predicted binding sites for the N-terminal matchstick region using the neural network DeepSite (Jiménez et al., 2017), which resulted in finding our identified binding sites for both hTIGIT and Gal-GalNAc with the highest prediction scores (SI Fig. 9e).

In summary, the Fap2/hTIGIT co-structure reveals important mechanistic details in the functional interaction of Fn with NK cells. First, co-binding of both Fap2 receptors is generally possible because the interfaces are spatially distant enough to avoid overlapping. Second, the location of the interaction interface at the very distant end of the long adhesin molecule, ∼50 nm away from Fn itself, with no further interaction sites identified at the flanks of the Fap2 rod, raises the question why Fn requires such a long adhesin protein to adhere to host cells. A possible hypothesis is that the long Fap2 rod acts as spacer to keep active NK cells on safe distance to Fn itself, but even more to tumors invaded by Fn, and deactivates the immune cells through hTIGIT interaction and subsequent signaling.

## Discussion

Here, we report the recombinant production, and the functional and structural analysis of the infection- and cancer-relevant fusobacterial autotransporter adhesin Fap2. The functionally relevant part of its ECD alone is already 338 kDa large, posing a great challenge for production in *E. coli*, which is by far the most widely used and established protein expression system. Only very few proteins above 2500 residues are reported in the protein data bank (PDB) with *E. coli* as expression system, i.e., Tc toxins (Gatsogiannis et al., 2018), Talin1 (Dedden et al., 2019), *M. tuberculosis* fatty acid synthase (Ciccarelli et al., 2013), *Clostridioides difficile* Toxin A (Chen et al., 2022). Fn Fap2-ECD in this work is, to our knowledge, the first membrane-associated protein of such a large size that has been produced in *E. coli* using the AIDA autotransporter system. This has previously only been used to display much smaller “passenger domains” on the *E. coli* surface (Lattemann et al., 2000; Jarmander et al., 2012), while our design proves its general applicability for the preparative production of surface proteins independent of their size and behavior in the *E. coli* cytoplasm.

Binding experiments to Gal-GalNAc-Thr and hTIGIT proved functionality of recombinantly produced Fap2-ECD. Using indirect binding competition experiments, we show that our recombinantly produced Fap2-ECD exhibits affinity to Gal-GalNAc-Thr that is at least as high as described for WT Fap2. For Fap2-ECD and hTIGIT, our measured K_D_ of ∼580 nM is the first reported quantitative value of this interaction. When compared to the affinities of other autotransporter adhesins with their receptors, e.g., *Haemophilus influenzae* autotransporter lipoprotein P4 with fibronectin and laminin (K_D_ about 10 nM) (Su et al., 2016), *E. coli* UpaB with fibronectin (K_D_ of 45 nM) (Paxman et al., 2019), or *Yersinia enterocolitica* YadA with collagen (K_D_ of 170 nM) (Leo et al., 2008), the Fap2/hTIGIT affinity appears weak. This is not unexpected when considering the binding of Fap2 to hTIGIT in the context of functional interaction with NK cells and tumor cells. It is the association to Gal-GalNAc that allows colonization of tumors (Abed et al., 2016; Parhi et al., 2020) and placenta (Parhi et al., 2022), probably through multivalent interactions due to low affinity of individual protein-glycan interactions (Cohen, 2015). A high density of Gal-GalNAc on the surface of tumor cells and a high expression level of Fap2 in Fn create the ideal conditions for this. In contrast, the physiological purpose of the Fap2/hTIGIT interaction is to deactivate NK cells and effector T-cells by inhibitory signaling through hTIGIT activation after Fap2 binding (Ge et al., 2021). Thereby, transient interaction is likely sufficient to initiate signaling and probably even favorable if hTIGIT binding interferes with Gal-GalNAc binding. Notably, the interaction of hTIGIT to Nectin-2 is an order of magnitude weaker and still sufficient to initiate hTIGIT signaling (Deuss et al., 2017; Lozano et al., 2020) with immune cell deactivation as consequence. Although our data indicate different binding sites, the possibility of interference cannot be ruled out due to a potential involvement of hTIGIT glycans.

Molecular docking and MD simulations identified Fap2 residues participating in the interaction with Gal-GalNAc and hTIGIT. *In silico* analysis of Fap2/Gal-GalNAc binding indicates the possibility for ambiguity of the binding pit towards further glycans. This finding is not surprising, as previous studies have demonstrated that the binding and invasion of CRC cells by Fn is affected by the presence of galactose-containing sugars (Casasanta et al., 2020). The Fap2-hTIGIT interaction includes, among multiple stable and consistent hydrogen bonds, a stacking-like interaction between a Pro and Trp residue which has been found to be stable and to play an important role in protein-protein interactions (Biedermannova et al., 2008; Zondlo, 2013). The *in silico* analysis of the Fap2-hTIGIT interaction complements the low resolution cryo-EM data and underlines the relevance of our integrated approach.

Taken together, our results suggest a unique mechanism of Fap2 interaction with both receptors, that takes the long shape of Fap2 and the hTIGIT binding site at its membrane-distal end into account. We propose that Fn applies Fap2 as molecular “spear” to deactivate tumor-invading immune cells already at the maximum possible distance through transient hTIGIT interaction and subsequent signaling (Fig. 5). At the same time, Fn is tightly associated to CRC cells in tumors via multiple Fap2/Gal-GalNAc interactions at the back side of the Fap2 tip, analogous to “grappling hooks”. This is enabled by high levels of Fap2 found on the Fn surface, as evident in Fig. 2a. Specific inhibition of either of these Fap2-mediated interactions is likely to have a tremendous impact on Fn tumor colonization and is thus a promising future strategy to combat CRC, particularly metastasis and chemotherapeutic resistance.

## Methods

### Source of plasmids and genes for design of expression constructs

The *E. coli* expression vector pAIDA1 was a gift from Gen Larsson (Addgene plasmid # 79180; http://n2t.net/addgene:79180; RRID:Addgene_79180) (Jarmander et al., 2012). The *fap2* gene sequence from *Fusobacterium nucleatum subsp. nucleatum* ATCC 23726 (FusoPortal Gene 2068 (Sanders et al., 2018)) was codon-optimized for expression in *E. coli* and purchased from GenScript. The hTIGIT gene sequence (Uniprot ID: Q495A1) was purchased from GenScript and cloned in a modified pEG BacMam vector with C-terminal eGFP (Vinayagam et al., 2018). pGEX4T3-GST-Tev was a gift from Robert Sobol (Addgene plasmid # 177142; http://n2t.net/addgene:177142; RRID:Addgene_177142) (Fang et al., 2019) and has been modified to replace the thrombin cleavage site by a HRV3C cleavage site.

### Cloning of expression plasmids for Fap2 and hTIGIT

The synthetic gene for the Fap2-ECD (residues 42 - 3271) from *Fusobacterium nucleatum subsp. nucleatum* ATCC 23726 was cloned into the pAIDA1 backbone in frame with the pAIDA1 signal peptide. A 8x His-tag was cloned between the C-terminal end of Fap2-ECD and the TEV cleavage site of pAIDA1 with 3 amino acid linkers before (GSA) and after (SAG) the His-tag, resulting in pAIDA_TEV_Fap2EC (Fig. 1d).

The ECD of hTIGIT including signal peptide (residues 1 - 141) was cloned into the pEG BacMam backbone downstream of the CMV promoter in fusion with a HRV3C site and a C-terminal eGFP with a GGS linker before eGFP. A Twin-Strep-tag was attached to the C-terminus of eGFP after a 5 amino acid linker (SGRMA).

All gene insertions were carried out using the NEBuilder HiFi DNA Assembly kit (New England Biolabs).

### Expression and purification of Fap2-ECD

BL21(DE3)star *E. coli* cells were transformed with pAIDA_TEV_Fap2EC, a positive colony was inoculated into 5 ml LB media and grown overnight at 37 °C. 1 ml aliquots were frozen at -80 °C in 25% sterile glycerol for cryopreservation. A 100 ml preculture was inoculated from the cryostock and grown at 37 °C and 180 rpm overnight. The *E. coli* cells were then pelleted at 3300 x g for 10 min and the supernatant was removed. 4 L of M9 minimal media supplemented with 5 mg/ml casamino acids, 100 µg/ml tryptophan and 35 µg/ml chloramphenicol were inoculated to an OD_600_ of ∼0.2 and incubated at 37 °C and 180 rpm. When an OD600 of ∼0.2 was reached, the temperature was lowered to 15 °C and expression was induced at an OD_600_ of ∼0.9 with 0.5 mM isopropyl-β-D-1-thiogalactopyranoside for subsequent overnight expression. Cells were harvested by centrifugation at 3300 x g for 9 min and resuspended in TEV cleavage buffer (20 mM TrisHCl pH 7.5, 200 mM NaCl, 10% glycerol, 0.5 mM EDTA, 1 mM DTT, 1 mM PMSF). GST tagged TEV protease (GST-TEV) was added to a concentration of ∼70 µg/ml and the cell suspension was incubated for 4 h at 6 °C on a rotary incubator. The released Fap2-ECD was then separated from the cells by centrifugation at 50,000 x g for 45 min. 10 mM MgCl_2_, 40 mM imidazole and 200 mM NaCl was added before loading on a self-packed 3 ml Ni Sepharose™ 6 Fast Flow (VWR) column using ÄKTA Pure FPLC system (Cytiva) at 4 °C. The bound protein was washed with 150 ml wash buffer (20 mM TrisHCl pH 7.5, 300 mM NaCl, 10% glycerol) and eluted with 36 ml elution buffer (20 mM TrisHCl pH 7.5, 300 mM NaCl, 10% glycerol, 300 mM imidazole). Collected fractions were analyzed by SDS-PAGE and Fap2-containing samples were pooled. Glutathione agarose resin (Cytiva) (1 ml) was used to remove remaining GST-TEV by batch incubation. Purified protein was either dialyzed into EM grid preparation buffer (20 mM TrisHCl pH 7.5, 200 mM NaCl) for vitrification or frozen in liquid nitrogen after adding glycerol to a concentration of 40% for storage at -80 °C.

### Expression and purification of hTIGIT-ECD

200 ml Expi293F™ cells were grown in Expi293™ Expression Medium (A14351-1, Gibco) in two separate 500 ml suspension cell culture flasks (100 ml each) at 37 °C, 90 rpm and 8% CO_2_. Transfection in each flask was performed at cell numbers between 1.5 and 2.0 x 10^6^ per mL and a minimum cell viability of 95%, using 1.5 µg plasmid per ml cell suspension. The plasmid solution was diluted with Opti-MEM (reduced serum free medium, 31985-047, Gibco) to a volume of 2.08 ml, and 225 µl PEI (Polysciences Europe) were added, followed by vortexing for 5 s. After incubation at 25 °C for 20 min, the DNA solution was added dropwise to the cell suspension. Incubation was continued at 37 °C, 90 rpm and 8% CO_2_.

After three days, the cells were harvested by centrifugation at 500 x g for 15 min at 4 °C and the pellet was frozen at -80 °C. The supernatant was collected and centrifuged for 30 min at 50.000 x g. Pierce™ protease inhibitor tablets (TFS) were added to the supernatant and then dialyzed against 4 L of buffer W (50 mM TrisHCl pH 8, 200 mM NaCl) overnight. The cell pellets were thawed on ice, resuspended in 10 ml buffer W (50 mM TrisHCl pH 8, 200 mM NaCl) and lysed by dounce homogenisation after addition of a Pierce™ protease inhibitor tablet (TFS). The lysate was centrifuged at 100.000 x g for 1 h at 4 °C and the cleared supernatant was pooled with the dialysed medium and applied to a Strep-Tactin® Sepharose® (IBA) gravity flow column containing 2 ml resin. The bound protein was washed with 25 column volumes (CVs) buffer W (50 mM TrisHCl pH 8, 200 mM NaCl) and eluted with 5 CVs Buffer E (50 mM TrisHCl pH 8, 200 mM NaCl, 2.5 mM Desthiobiotin). Elution fractions were analyzed for GFP emission signal, pooled accordingly and concentrated using Amicon®Ultra-4 centrifugal filter unit (Merck Millipore) to a final volume of 1 ml. Size exclusion chromatography was then performed with buffer W (50 mM TrisHCl pH 8, 200 mM NaCl) using a Superdex 200 Increase 10/300 GL column to separate free GFP from hTIGIT-ECD. Fractions containing hTIGIT-ECD were pooled from two runs and directly frozen in liquid nitrogen after addition of 40% glycerol for storage at -80 °C.

We also produced non-glycosylated hTIGIT in *E. coli* and purified it from inclusion bodies as described previously (Stengel et al., 2012).

### Cell binding assay of Fap2-expressing *E. coli* to cancer cells

For quantitative cell binding experiments with HT-29 cells (ATCC), µ-slide 8 well glass bottom plates (Ibidi) were coated by adding 0.1 mg/ml poly-L-lysine to each well for 1 h at 37 °C with subsequent washing with sterile H_2_O and drying for 2 h at 25 °C. 120,000 HT-29 cells were seeded in 300 µl DMEM/F12 + 10% FBS (PAN-Biotech) per well and incubated at 37 °C and 5% CO_2_ for 48 h.

The pAIDA_TEV_Fap2EC plasmid was used to express Fap2 on the outer membrane in *E. coli* as described above. *E. coli* cells were harvested by centrifugation at 3200 x g for 5 min, washed three times with PBS pH 7.4 and adjusted to an OD_600_ of 5.0 in 700 µl. Plasma membranes were stained by adding 3.5 µl CellBrite® Fix 555 (Biotium) for 30 min at 25 °C in the dark. The cells were harvested by centrifugation at 3200 x g for 5 min, washed once with PBS and resuspended to an OD_600_ of 4.0. For assays in the presence of GalNAc, stained *E. coli* (OD_600_ of 2.0) were incubated with 300 mM GalNAc for 30 min at 25 °C in the dark and diluted 1:1 with PBS upon adding 80 µl of the suspension to HT-29 cells. If no sugar was used, stained *E. coli* were adjusted to an OD_600_ of 1 and directly added to HT-29 cells. As a control, *E. coli* cells were transformed using the empty pAIDA plasmid and treated similarly to the Fap2-expressing *E. coli*. After addition of *E. coli* to the HT-29 cells, they were incubated at 37 °C and 5% CO_2_ for 1 h. The cells were then washed twice with PBS, fixed with paraformaldehyde (Roth) for 15 minutes at 25 °C and washed with PBS containing 1% Tween-20 before staining with 150 µl Phalloidin-iFluor 488 (Abcam) for 20 min at 25 °C. After washing twice with PBS, the cells were imaged using a Zeiss LSM780 confocal microscope. The number of *E. coli* attached per HT-29 cell was counted and analyzed statistically using the Games-Howell post-hoc test from the ggstatsplot package (Patil, 2021) in R.

Qualitative attachment assays of Fap2-expressing *E. coli* to Caco-2 (ATCC) cells were conducted in a similar way as described for HT-29 cells with following deviations: The assay was performed in 24-well cell culture plates (Sarstedt) with a total volume of 500 µl per well. Caco-2 cells were not stained and imaged in bright field.

### Labeling of Gal-GalNAc-Thr with AF 488-NHS and binding to Fap2-expressing *E. coli*

Gal-GalNAc-Thr (4.13 µmol) was dissolved in 200 mM bicarbonate buffer (1 ml, pH 8.3) and labeled with AF 488-NHS (Lumiprobe, Cat.Nr.: 21820) dissolved in 10 µl DMSO in a 1:1.5 ratio for 1 h at 25 °C. Excess AF 488 was removed after labeling by high-performance liquid chromatography (HPLC) using a preparative column (Reprospher 100 C18, 10 μm: 50 x 30 mm at 20 mL/min flow rate, linear gradient from H2O/MeCN + 0.1 trifluoracetic acid from 10 to 90% over 60 min) on a 1260 Infinity II LC System (Agilent). Peaks were collected manually according to the peak height at 480 nm. Separation was confirmed by mass detection in collected fractions using a single quadrupole LC/MSD system (Agilent) and fractions containing Gal-GalNAc-Thr-AF488 (0.45 µmol, yield: 10.9 %, calc. for C39H43N4O23S2^-^ [M-H]^-^: 999.18, found: 999.2, negative mode) were pooled and lyophilized for storage at -20 °C.

Fap2-expressing *E. coli* were prepared as described above. *E. coli* transformed with an empty pAIDA plasmid were used as a control. Gal-GalNAc-Thr-AF488 was dissolved in PBS pH 7.4 (200 µM) and added to the bacteria. The final concentration range was from 0 to 100 µM in a total volume of 400 µl with an *E. coli* density of OD_600_ of 1.5. Bacteria were incubated for 1 h on ice, followed by 30 min at 37 °C and then pelleted for 7 min at 3.300 x g. After washing the pellet with PBS, it was resuspended in 160 µl lysis buffer (40 µl BugBuster (Merck Millipore) + 120 µl PBS).

0.5 µl Benzonase® (MoBiTec) was added and *E. coli* were lysed for 20 min at 37 °C, 600 rpm. The lysate was cleared by centrifugation at 16.000 x g for 20 min at 25 °C and fluorescence signal at 519 nm (excitation wavelength: 459 nm) was detected in a 96-well flat bottom microplate (Corning) with a Safire microplate reader (Tecan). Fluorescence detected in control *E. coli* was subtracted from that of induced Fap2-expressing *E. coli*. The resulting difference was plotted in relation to the Gal-GalNAc-Thr concentration.

### Pulldown assay of Fap2-ECD and hTIGIT-ECD

Purified Fap2-ECD and hTIGIT-ECD were thawed on ice from frozen aliquots (500 µl each). Fap2-ECD was immobilized on 5% Ni-NTA magnetic beads for 2.5 h at 8 °C in a rotary shaker and washed three times with wash buffer (20 mM TrisHCl pH 7.5, 150 NaCl, 5 mM Imidazole). Thawed hTIGIT-ECD was added to the magnetic beads and incubated for 2.5 h at 25 °C in a rotary shaker. Unbound hTIGIT-ECD was removed by washing three times with wash buffer and the complex was then eluted from the beads with elution buffer (20 mM TrisHCl pH 7.5, 150 NaCl, 500 mM Imidazole).

### Surface plasmon resonance of Fap2-ECD and hTIGIT-ECD

Binding kinetics were determined using a Biacore™ 3000 surface plasmon resonance system (Cytiva). hTIGIT-ECD in fusion with a C-terminal Fc tag (Acro Biosystems, Cat. No. TIT-H5254) was immobilized on CM5 sensor chips (GE Healthcare) in 10 mM sodium acetate, pH 4.8, using a mixture of 0.2 M 1-ethyl-3-(3-dimethylaminopropyl)-carbo-diimide (EDC) and 50 mM N-hydroxysuccinimide (NHS) supplied by the manufacturer. The injection of 10 µg/ml hTIGIT-Fc for 7 min with a flow of 10 µl/min resulted in a final responsive unit (RU) value of 3220. Excess reactive groups were deactivated using 1 M ethanolamine-HCL, pH 8.5. 50 µl Fap2-ECD was injected in different concentrations ranging from 52,5 to 368 nM and the response difference was recorded. All measurements were carried out in 25 mM HEPES-NaOH pH 7.5, 50 mM NaCl, and 0.005% Tween-20. Analyses were performed at 25 °C with a flow rate of 20 μl/min for 2.5 min and dissociation for 3 min. After each run, surfaces were regenerated with 30 µl 50 mM NaOH, 1 M NaCl at a flow rate of 20 μl/min for 1.5 min. The BIAevaluation software v 4.1.1 (Pharmacia Biosensor) was used to calculate association and dissociation rate constants by 1:1 Langmuir fitting of the primary sensorgram data.

### AlphaFold2 prediction of Fap2 ATCC23726, Fap2 ATCC25586 and RadD ATCC23726

A local installation of AlphaFold2 (Jumper et al., 2021) was used to predict the structures of Fap2 (FusoPortal Gene 2068) and RadD (FusoPortal Gene 32) from *Fusobacterium nucleatum subsp*.

*nucleatum* ATCC 23726 and Fap2 from *Fusobacterium nucleatum subsp. nucleatum* ATCC 25586 (FusoPortal Gene 1976). Due to the size of these proteins, predictions were carried out for five separate parts with overlapping regions of at least 100 amino acids, covalently assembled in ChimeraX and energy minimized using ISOLDE (Croll, 2018).

### Negative stain EM

4 μl of either purified, detergent solubilized Fap2 from Fn ATCC 25586 or recombinantly purified Fap2-ECD at a concentration of 0.05 mg/ml was applied to a freshly glow-discharged 300-mesh carbon-coated copper grid. After an incubation of 1 min, the sample was blotted with Whatman filter paper, washed twice by application of 4 µl Milli-Q water with subsequent blotting and stained with 4 µl 2% uranyl acetate solution for 1 min. The staining solution was removed with Whatman filter paper and grids were air-dried for at least 5 min at room temperature. Grids were imaged on a Talos L120 TEM (TFS) using a Ceta-16M CCD detector at 73,000x magnification.

### Cryo-EM sample preparation and data collection of Fap2-ECD and Fap2-ECD/hTIGIT-ECD

Purified Fap2-ECD at a concentration of 0.1 mg/ml was vitrified on Quantifoil R 2/1 Cu 300 mesh grids using a Vitrobot Mark IV (TFS) by applying 4 µl of sample twice with subsequent blotting for 2.0 s and 4.0 s, respectively, with a blotting force of -1, and 100% humidity at 12 °C.

Purified Fap2-ECD in complex with hTIGIT was eluted from magnetic beads and used for vitrification on Quantifoil R 2/1 Cu 300 mesh grids with a Vitrobot Mark IV (TFS). 4 µl of sample was applied 3 times with subsequent manual blotting steps using Whatman paper from the back side of the grid. Grids were vitrified after pipetting 4 µl buffer (20 mM Tris, 150 mM NaCl) onto the grid to reduce imidazole concentration. Blotting was done for 3.0 s, with a blot force of -1 and a humidity of 100% at 12 °C.

Micrographs were acquired on a Titan Krios G3i microscope (TFS) operated at 300 kV equipped with a K3 direct electron detector and Bioquantum energy filter (Gatan, Digital Micrograph version 3.32.2403.0) running in CDS superresolution counting mode at a slit width of 20 eV and at a nominal magnification of 81,000x, giving a calibrated pixel size of 0.53 Å/px. For Fap2-ECD, movies were recorded for 2.0 s accumulating a total electron dose of 59.8 e^−^/Å^2^ fractionated into 53 frames. For Fap2-ECD/hTIGIT-ECD, movies were recorded for 2.0 s accumulating a total electron dose of 44.6 e^−^/Å^2^ fractionated into 38 frames. EPU 2.12 was utilized for automated data acquisition using nominal defocus values between -1.5 and -2.6 µm for Fap2-ECD, and between -1.0 and -2.5 µm for Fap2/hTIGIT-ECD, respectively.

### Cryo-EM data analysis and modeling of Fap2-ECD

Processing of the cryo-EM data was performed in CryoSPARC (Punjani et al., 2017) and is outlined in SI Fig. 4. 12,663 movies were collected, aligned using Patch Motion correction and CTF was determined with patch CTF estimation. After sorting out bad images, 12,391 micrographs were kept and processed further. A small subset of 59 micrographs was used to manually pick 474 particles to train Topaz (Bepler et al., 2019). 330,980 particles were then picked using the trained Topaz model, extracted (7x binned), and subjected to 2D classification (30 online-EM iterations). This process was iterated by repeating twice, using 30,000 and 52,444 particles of selected 2D classes to train new Topaz models. The final round of 2D classification yielded 321,160 particles after extraction from micrographs without binning (1.06 Å/px) which were used for an ab-initio reconstruction. Subsequent homogenous and non-uniform refinement followed, the latter using the automatically generated static mask from the homogenous refinement. Particles were then subjected to two further rounds of 2D classification (30 online-EM iterations each), leaving 265,956 particles. These were used for non-uniform refinement of the previously obtained volume, which resulted in a map with a resolution of 4.7 Å according to the gold-standard Fourier shell correlation (FSC) criterion (SI Fig. 3b, SI Table 1).

To optimize resolution in the N-terminal matchstick region, the following steps were carried out: The volume from the first non-uniform refinement was used to fit the AlphaFold2 prediction of Fap2 from Fn ATCC 23726 (residues 42 - 2698) into the density. The model was automatically fitted using the flexible molecular dynamics fitting software Namdinator (Kidmose et al., 2019) and manually adjusted in Coot (Emsley et al., 2010). The Fap2 model was then assembled in ChimeraX (Pettersen et al., 2021) with an AlphaFold2 prediction to include a C-terminal section of the Fap2-ECD construct that could not be fitted into the density. The resulting model of Fap2-ECD (42 - 3271) was then used to generate an artificial map in ChimeraX with a resolution of 5 Å, which was applied as a template for a non-uniform refinement, using the cleaned-up particles from the final 2D classification. A mask (Dilation radius 8, soft padding width 15) covering the N-terminal end was generated in ChimeraX, followed by 3D classification to find particles with good alignment on the matchstick region. 160,806 particles were retained and subjected to particle subtraction of the C-terminal region using a mask (Dilation radius 10, soft padding width 20) generated in ChimeraX. 160,676 subtracted particles were used for local refinement, and the resulting volume and particles were re-aligned with the volume alignment tools job to shift the N-terminal matchstick region to the center of the box (new center: 350, 350, 417; box of 700 px). A local refinement job was carried out using the shifted particles, and particles connected to the resulting volume were re-extracted from micrographs with a box size of 384 px. A mask (Dilation radius 6, soft padding width 12) covering most of the N-terminal region was generated in ChimeraX, followed by local refinement and 3D classification using the subtracted particles. 151,825 particles were selected and a final local refinement resulted in a map with a global resolution of 4.4 Å, according to the gold-standard FSC criterion (SI Fig. 3f). DeepEMhancer (Sanchez-Garcia et al., 2021) was applied for map sharpening and an AlphaFold2 prediction of Fap2 (283 - 1817) was fitted using Namdinator (Fig. 1g,h). In addition, particles underlying the last refinement were imported in Relion (Scheres, 2012) using pyEM (Asarnow et al., 2019), refined and postprocessed, resulting in a map with 4.4 Å resolution according to the gold-standard FSC criterion.

### Cryo-EM data analysis of Fap2-ECD/hTIGIT-ECD

Processing of the cryo-EM data was performed in CryoSPARC and is outlined in SI Fig. 8f. Two data sets were acquired and combined, resulting in 15,658 patch CTF-estimated, motion-corrected micrographs after discarding low-quality images. Initially, 455 particles were picked manually on 49 micrographs to train Topaz. The trained model was used to extract 659,696 particles from micrographs with 3x binning, followed by 2D classification, and 41,675 particles were selected to train Topaz again. Subsequent picking resulted in 1,191,741 particles, which were extracted from micrographs (4x binning) and applied to two rounds of 2D classification (50 online-EM iterations). An ab-initio reconstruction (5 classes) of the remaining 761,580 particles was used for an initial 3D sorting step, and 622,717 particles were retained for extraction (no binning) and two iterative rounds of 2D classification (50 online-EM iterations). 73,482 particles were selected for an ab-initio reconstruction, which was used for subsequent homogenous and non-uniform refinement, the latter using the mask from the homogenous refinement. The volume alignment tools job was applied to re-align particles to shift the N-terminal matchstick region to the center of the box (new center: 350, 350, 210; box of 700 px). The re-centered particles were used for an ab-initio reconstruction and further refined with subsequent homogenous and non-uniform refinement, with the mask generated in homogenous refinement. The resulting volume was used to create a mask in ChimeraX covering the matchstick region, with a dilation radius of 6 and soft padding width of 20. The N-terminal focus mask was applied for two independent 3D classifications in simple and input modes, respectively, both with “hard classification” enabled. Classes were selected based on the presence of extra density at the N-terminus, yielding 45,207 particles. These were further processed using non-uniform refinement to a final global resolution of 6.0 Å according to the gold-standard Fourier shell correlation (FSC) criterion (SI Fig. 8c, SI Table 1).

### Cultivation of Fn ATCC 25586

*F. nucleatum* ATCC 25586 (Leibniz Institute DSMZ, German Collection of Microorganisms and Cell Cultures) was grown at 37 °C under anaerobic conditions in reduced Schaedler broth (Sifin diagnostics). After overnight culture, to a McFarland standard 3, the bacteria were harvested at 3000 x g for 10 min and the supernatant was replaced with fresh medium. 5 ml of Jurkat cells (3.2 x 10^6^ cells/ml) were added to 45 ml of Fn culture and incubated overnight at 37 °C under anaerobic conditions. The cell suspension was harvested the next day at 3,000 x g for 10 min and frozen at -80 °C.

### Cryo-ET of Fn ATCC 25586 and tomogram reconstruction

A frozen sample of *Fusobacterium nucleatum* ATCC 25586 cells was thawed and diluted in 1x PBS pH 7.4 to an OD_600_ of ∼0.3. 4 µl of the bacterial suspension was mixed with 10 nm gold fiducials, added to a Quantifoil R 1.2/1.3 300 mesh grid and blotted for 3.0 s, with a blot force of 3 and humidity of 100% at 12 °C using a Vitrobot Mark IV (TFS). Imaging was performed on a FEI Titan Krios G3i microscope (TFS) operated at 300 kV equipped with a K3 direct electron detector and Bioquantum energy filter (Gatan, Digital Micrograph version 3.32.2403.0) running in CDS counting mode at a slit width of 20 eV, operated by SerialEM v4.0.4 (Mastronarde, 2003). A nominal magnification of 53,000x was used, which resulted in a calibrated pixel size of 1.68 Å/px. Tilt series were acquired using a dose-symmetric scheme (Hagen, Wan and Briggs, 2017), with a tilt axis angle of 84.4°, 3° angular increment and target defocus of -5.5 μm. Tilt series were recorded as movies of 10 frames in counting mode and a dose rate of 5.65 e−/Å^2^/sec resulting in a total dose of 160.7 e−/Å^2^ per tilt series. Frames were aligned using Warp/M (Tegunov and Cramer, 2019). Tomograms were reconstructed in etomo/IMOD (v.4.11.24) (“Computer Visualization of Three-Dimensional Image Data Using IMOD.,” no date), using ‘ctf phase flip’ for CTF correction and back projection with Simultaneous Iterative Reconstruction Technique (SIRT)-like filter equivalent to 15 iterations.

### Isolation of Fn adhesins from outer membrane

Frozen suspensions of *F. nucleatum* sp. ATCC 25586 were thawed, resuspended in lysis buffer (50 mM Tris-HCl pH 7.2, 300 mM NaCl, 10% glycerol), and lysed using a LM10 Microfluidizer (Microfluidics) at 15,000 psi and 4 °C, followed by removal of cell debries via centrifugation for 30 min at 50,000 x g and 4 °C. The membranes in the cleared lysate were pelleted for 2 h at 100,000 x g and 4 °C. Membrane pellets were homogenized in lysis buffer (50 mM Tris-HCl pH 7.2, 300 mM NaCl, 10% glycerol) using dounce homogenization, and 1% n-dodecyl β-D-maltoside (DDM) was added for solubilization overnight at 8 °C while shaking. Non-solubilized material was removed by centrifugation at 180,000 x g and 4 °C for 1h, and the supernatant was loaded on a Superose 6 10/300 column, using a buffer containing 50 mM Tris-HCl pH 7.2, 300 mM NaCl, 10% glycerol and 0.05% DDM. Collected fractions were analyzed by SDS-PAGE and negative stain EM.

### Peptide fingerprint of purified Fap2-ECD, and Fap2 and RadD from Fn ATCC 25586

Purified recombinant Fap2-ECD was further purified on a HiLoad 16/600 Superose 6 pg column (Cytiva) equilibrated in 50 mM Tris-HCl pH 7.4, 300 mM NaCl, 10% glycerol before MS analysis. Isolated outer membrane proteins from Fn ATCC 25586 were used after the Superose 6 (see above). Proteins were loaded on SDS-PAGE and subjected to in-gel digestion. For this, gel bands were excised, reduced with 5 mM DTT at 56 °C for 30 min and alkylated with 40 mM chloroacetamide at room temperature for 30 min in the dark. Protein digestion was carried out using trypsin at an enzyme-to-protein ratio of 1:100 (w/w) at 37 °C overnight. LC-MS analysis was performed using an UltiMate 3000 RSLC nano LC system coupled on-line to an Orbitrap Fusion mass spectrometer (Thermo Fisher Scientific). Reversed-phase separation was performed using a 50 cm analytical column (in-house packed with Poroshell 120 EC-C18, 2.7µm, Agilent Technologies) with a 120 min gradient. MS1 scans were performed in the Orbitrap using 120,000 resolution; MS2 scans were acquired in the ion trap with an AGC target of 10,000 and maximum injection time of 35 ms, charge state 2-4 enable for MS2. Data analysis including label free quantification was performed using MaxQuant (version 1.6.2.6 and 2.0.3.0) using the following parameters: MS ion mass tolerance: 4.5 ppm; MS2 ion mass tolerance: 0.5 Da; variable modification: Met oxidation, Acetyl (Protein N-term), Cys carbamidomethyl; protease: Trypsin (R,K); allowed number of missed-cleavages: 2, database: SwissProt and *E. coli* database (SwissProt_14dez, Escherichia_coli_2020) combined with the sequences Fap2-ECD from Fn ATCC 23726 and TEV protease, as well as Fap2 and RadD from Fn ATCC 25586; label free quantification; match between runs disabled. Results were reported at 1% false discovery rate at the protein level.

### Molecular docking of hTIGIT-ECD to Fap2-ECD

The Fap2-ECD model (residues 242 - 2698) was rigid-body fitted in the cryo-EM density map of the Fap2-ECD/hTIGIT-ECD co-structure using UCSF ChimeraX. Molecular docking of hTIGIT-ECD (residues 22 - 141, PDB ID 3UCR) was carried out with the HADDOCK2.2 (High Ambiguity Driven protein-protein DOCKing) web server (van Zundert et al., 2016). Amino acids forming the distal surface of the matchstick region (Fap2 residues 362-373, 533-544, 548-551, 594-595, and 720-721) were selected for the docking in accordance to the extra density observed for the Fap2-hTIGIT co-structure in comparison to the structure of Fap2 alone (Fig.4b,c). The cluster with the best docking results (SI Table 3) was further analyzed in MD simulation.

### Molecular docking of Gal-GalNAc-Thr to Fap2-ECD

Gal-GalNAc-Thr was docked to the AlphaFold2 predicted Fap2 model (339 - 910) using Autodock Vina (Eberhardt et al., 2021) in UCSF Chimera (V1.17.1) (Pettersen et al., 2004). Protonation of Fap2 was set to charge Asp, Glu, and Lys. The search frame was set to include the matchstick region (D339 – N910) at the distal end of Fap2. The best docking pose gave a free energy score of -6.1 kcal/mol (SI Fig.9a). To confirm the relevance of the docking, it was performed in three replicates searching for ten states each on a larger N-terminal area of Fap2 (312 - 1381) and compared to docking results obtained for the N-terminal region of RadD (474 - 1274). The docking results were visualized in ChimeraX and colored according to the docking score (SI Fig. 9b,c).

### MD simulation

The best molecular docking results were analyzed by MD simulations using the GROMACS 2022 package (Abraham et al., 2015) with the CHARMM36 force field, which is compatible with proteins and glycoproteins (Huang and MacKerell Jr, 2013), and the standard water model CHARMM-modified TIP3P (Boonstra, Onck and van der Giessen, 2016).

For the simulation of the Fap2/Gal-GalNAc-Thr complex, the docked structure was split into receptor and ligand structure for further processing. The receptor topology was generated using the *pdb2gmx* program with interactive selection of NH3^+^ and COO^-^ termini. This manual selection of protein-specific termini is necessary due to the N-terminal methionine residue for which *pdb2gmx* selects the incompatible residue type t by default. The docked ligand structure was hydrogenated and converted to .mol2 format using UCSF Chimera (Pettersen et al., 2004). The ligand topology was generated using the online tool SwissParam (Bugnon et al., 2023). The [atomtypes] and [pairtypes] entries from the resulting .itp file were transferred to the resulting .prm ligand file, which was then converted to the GROMACS .gro format using *gmx editconf.* Finally, the receptor and ligand .gro files were merged and the ligand was added to the topology file. For the simulation of the Fap2-hTIGIT complex, the topology generated directly of the complete docked structure.

The complex was immersed in a dodecahedral box with edge spacing of 1 nm. The box was filled with spc216 water molecules and sodium ions to neutral pH using *gmx editconf*, *solvate*, *grompp* and *genion*. The system was energy minimized using the steepest integrator and Verlet cutoff scheme. NVT equilibration was performed using the md leap-frog integrator in 50000 2 fs steps with V-scale temperature coupling between protein and non-protein groups with a 300 K reference temperature. NPT equilibration was performed using Parrinello-Rahman pressure coupling. A Verlet cutoff scheme was used for van der Waals interactions and the Ewald particle mesh for long-range electrostatic interactions. No distance constraints were applied. The production simulation was performed using the same parameters for 50 ns.

The resulting trajectories were fixed and centered with *gmx trjconv* and analyzed with the vmd hbonds plugin 1.2 (Humphrey, Dalke and Schulten, 1996) and the GROMACS-implemented RMSF tool (Abraham et al., 2015) to detect movements of the ligand in the binding interface. Hydrogen bond analysis was performed by selecting all receptor residues within 5 Å of the ligand in the selected docking state as selection 1 and the ligand as selection 2. A distance cutoff of 3.5 nm and an angle cutoff of 35° were used, and the results of multiple replicate trajectories were statistically analyzed using a custom Python script. The most consistent interaction partners were visually inspected in vmd. Representative trajectories were rendered using the vmd Movie Maker plugin (Humphrey, Dalke and Schulten, 1996) combined with FFmpeg (“Converting Video Formats with FFmpeg | Linux Journal,” no date).

### Binding site prediction of Fap2-ECD matchstick region

The neural network based binding site predictor DeepSite (Jiménez et al., 2017) was utilized to predict binding sites of the Fap2 matchstick region. An AlphaFold2 prediction of Fap2 (339 - 910) was used for analysis on the playmolecule web-based server (www.playmolecule.org) and results were visualized with ChimeraX, coloring the predicted sites according to their score.

## Data and code availability

Cryo-EM densities of Fap2 and Fap2/hTIGIT have been deposited at the EMDB under accession numbers … and …, respectively. The source data underlying Figures and Supplementary Figures are provided as a Source Data file. The data sets and scripts generated in this work are available from the corresponding author on request.

## Acknowledgements

We thank Uwe Fink, Alina Roderer (Leibniz FMP), and Julia Schmidt (Charité - Universitätsmedizin Berlin) for excellent technical support, and Marianne Schoenfelder (TU Berlin) for assistance in TIGIT purification. We thank Dr. Oxana Krylova (Biophysics facility, Leibniz FMP) for support with recording and analyzing SPR data. We thank Dr. Johannes Broichhagen, Dr. Blaise Gatin-Fraudet, and Kilian Rossmann (ChemBioProbes, Leibniz FMP) for their help with Gal-GalNAc-Thr labeling and purification, and Dr. Peter Schmieder and Nils Trieloff (NMR Core facility, Leibniz FMP) for NMR analysis of labeled Gal-GalNAc-Thr. We acknowledge peptide fingerprint analysis carried out and analyzed by Heike Stephanowitz (Mass Spectrometry Core facility, Leibniz FMP). We acknowledge training for and access to the Zeiss confocal microscope by Marie Bieck and Martin Lehmann (Cellular imaging facility, Leibniz FMP). We acknowledge access to electron microscopic equipment at the Core Facility for cryo-Electron Microscopy (CFcryoEM) of the Charité - Universitätsmedizin Berlin supported by DFG (INST 335/588-1 FUGG) for cryo-EM and cryo-ET data collection, and we thank Dr. Christoph Diebolder (CFCryoEM) for support in cryo-ET data analysis.

## Author contribution

FS designed expression constructs, carried out and evaluated Fap2 production, cryo-EM, cryo-ET, and functional assays. GLM carried out and analyzed MD simulations. KM carried out and analyzed cell adhesion assays. TS recorded cryo-EM and cryo-ET data, including data preprocessing. AM supervised and JK carried out and optimized Fn cultivation. DR designed and supervised research and carried out docking and modeling, together with FS. DR, FS, and GLM prepared figures, movies, and wrote the manuscript with contributions from all authors.

## References

1. Abed, J. et al. (2016) “Fap2 Mediates Fusobacterium nucleatum Colorectal Adenocarcinoma Enrichment by Binding to Tumor-Expressed Gal-GalNAc.”

2. Abraham, M. J. et al. (2015) “GROMACS: High performance molecular simulations through multi-level parallelism from laptops to supercomputers,” SoftwareX, 1–2, pp. 19–25. doi: 10.1016/j.softx.2015.06.001.

3. Ai, D. et al. (2019) “Identifying gut microbiota associated with colorectal cancer using a zero-inflated lognormal model.”

4. Arnold, M. et al. (2017) “Global patterns and trends in colorectal cancer incidence and mortality,” Gut, 66(4), pp. 683–691. doi: 10.1136/gutjnl-2015-310912.

5. Arnold, M. et al. (2020) “Global Burden of 5 Major Types of Gastrointestinal Cancer,” Gastroenterology, 159(1), pp. 335–349.e15. doi: 10.1053/j.gastro.2020.02.068.

6. Bepler, T. et al. (2019) “Positive-unlabeled convolutional neural networks for particle picking in cryo-electron micrographs,” Nat Methods, 16(11), pp. 1153–1160. doi: 10.1038/s41592-019-0575-8.

7. Biedermannova, L. et al. (2008) “Another role of proline: stabilization interactions in proteins and protein complexes concerning proline and tryptophane,” Physical chemistry chemical physics: PCCP, 10(42), pp. 6350–6359. doi: 10.1039/b805087b.

8. Boonstra, S., Onck, P. R. and van der Giessen, E. (2016) “CHARMM TIP3P Water Model Suppresses Peptide Folding by Solvating the Unfolded State,” The Journal of Physical Chemistry B, 120(15), pp. 3692–3698. doi: 10.1021/acs.jpcb.6b01316.

9. Brennan, C. A. and Garrett, W. S. (2019) Fusobacterium nucleatum — symbiont, opportunist and oncobacterium, Nature Reviews Microbiology.

10. Bugnon, M. et al. (2023) “SwissParam 2023: A Modern Web-Based Tool for Efficient Small Molecule Parametrization,” Journal of Chemical Information and Modeling, 63(21), pp. 6469–6475. doi: 10.1021/acs.jcim.3c01053.

11. Casasanta, M. A. et al. (2020) “Fusobacterium nucleatum host-cell binding and invasion induces IL-8 and CXCL1 secretion that drives colorectal cancer cell migration,” Sci Signal, 13(641). doi: 10.1126/scisignal.aba9157.

12. Castellarin, M. et al. (2012) “Fusobacterium nucleatum infection is prevalent in human colorectal carcinoma.”

13. Chen, B. et al. (2022) “Structure and conformational dynamics of Clostridioides difficile toxin A,” Life Science Alliance, 5(6), p. e202201383. doi: 10.26508/lsa.202201383.

14. Ciccarelli, L. et al. (2013) “Structure and Conformational Variability of the Mycobacterium tuberculosis Fatty Acid Synthase Multienzyme Complex,” Structure, 21(7), pp. 1251–1257. doi: 10.1016/j.str.2013.04.023.

15. Cohen, M. (2015) “Notable aspects of glycan-protein interactions,” Biomolecules, 5, pp. 2056–2072. doi: 10.3390/biom5032056.

16. “Computer Visualization of Three-Dimensional Image Data Using IMOD.” (no date). Available at: http://bio3d.colorado.edu/imod/paper/ (Accessed: January 22, 2024).

17. “Converting Video Formats with FFmpeg | Linux Journal” (no date). Available at: https://www.linuxjournal.com/article/8517 (Accessed: January 24, 2024).

18. Coppenhagen-Glazer, S. et al. (2015) “Fap2 of Fusobacterium nucleatum is a galactose-inhibitable adhesin involved in coaggregation, cell adhesion, and preterm birth.”

19. Croll, T. I. (2018) “ISOLDE: a physically realistic environment for model building into low-resolution electron-density maps,” Acta Crystallogr D Struct Biol, 74(Pt 6), pp. 519–530. doi: 10.1107/S2059798318002425.

20. Dai, Z. et al. (2018) “Multi-cohort analysis of colorectal cancer metagenome identified altered bacteria across populations and universal bacterial markers.” NLM (Medline).

21. Dedden, D. et al. (2019) “The Architecture of Talin1 Reveals an Autoinhibition Mechanism,” Cell, 179(1), pp. 120–131.e13. doi: 10.1016/j.cell.2019.08.034.

22. Deuss, F. A. et al. (2017) “Recognition of nectin-2 by the natural killer cell receptor T cell immunoglobulin and ITIM domain (TIGIT),” The Journal of Biological Chemistry, 292(27), pp. 11413–11422. doi: 10.1074/jbc.M117.786483.

23. Eberhardt, J. et al. (2021) “AutoDock Vina 1.2.0: New Docking Methods, Expanded Force Field, and Python Bindings,” Journal of Chemical Information and Modeling, 61(8), pp. 3891–3898. doi: 10.1021/acs.jcim.1c00203.

24. Emsley, P. et al. (2010) “Features and development of Coot,” Acta Crystallographica Section D: Biological Crystallography, 66(4), pp. 486–501. doi: 10.1107/S0907444910007493.

25. Fang, Q. et al. (2019) “Stability and sub-cellular localization of DNA polymerase β is regulated by interactions with NQO1 and XRCC1 in response to oxidative stress,” Nucleic Acids Research, 47(12), pp. 6269–6286. doi: 10.1093/nar/gkz293.

26. Gatsogiannis, C. et al. (2018) “Tc toxin activation requires unfolding and refolding of a β-propeller,” Nature, 563, pp. 209–233. doi: 10.1038/s41586-018-0556-6.

27. Ge, Z. et al. (2021) “TIGIT, the Next Step Towards Successful Combination Immune Checkpoint Therapy in Cancer,” Front Immunol, 12, p. 699895. doi: 10.3389/fimmu.2021.699895.

28. GlycoSHIELD: a versatile pipeline to assess glycan impact on protein structures | bioRxiv (no date). Available at: https://www.biorxiv.org/content/10.1101/2021.08.04.455134v3 (Accessed: January 17, 2024).

29. Guo, P. et al. (2020) “FadA promotes DNA damage and progression of Fusobacterium nucleatum-induced colorectal cancer through up-regulation of chk2.” BioMed Central Ltd.

30. Gur, C. et al. (2015) “Binding of the Fap2 protein of fusobacterium nucleatum to human inhibitory receptor TIGIT protects tumors from immune cell attack.”

31. Hagen, W. J. H., Wan, W. and Briggs, J. A. G. (2017) “Implementation of a cryo-electron tomography tilt-scheme optimized for high resolution subtomogram averaging.” Academic Press Inc.

32. Han, Y. W. et al. (2004) “Fusobacterium nucleatum Induces Premature and Term Stillbirths in Pregnant Mice: Implication of Oral Bacteria in Preterm Birth.” American Society for Microbiology Journals.

33. Huang, J. and MacKerell Jr, A. D. (2013) “CHARMM36 all-atom additive protein force field: Validation based on comparison to NMR data,” Journal of Computational Chemistry, 34(25), pp. 2135–2145. doi: 10.1002/jcc.23354.

34. Humphrey, W., Dalke, A. and Schulten, K. (1996) “VMD: Visual molecular dynamics,” Journal of Molecular Graphics, 14(1), pp. 33–38. doi: 10.1016/0263-7855(96)00018-5.

35. Ito, M. et al. (2015) “Association of Fusobacterium nucleatum with clinical and molecular features in colorectal serrated pathway.” Wiley-Liss Inc.

36. Jarmander, J. et al. (2012) “A dual tag system for facilitated detection of surface expressed proteins in Escherichia coli,” Microb Cell Fact, 11, p. 118. doi: 10.1186/1475-2859-11-118.

37. Jiménez, J. et al. (2017) “DeepSite: protein-binding site predictor using 3D-convolutional neural networks,” Bioinformatics, 33(19), pp. 3036–3042. doi: 10.1093/bioinformatics/btx350.

38. Jones, P. et al. (2014) “InterProScan 5: genome-scale protein function classification,” Bioinformatics, 30(9), pp. 1236– 1240. doi: 10.1093/bioinformatics/btu031.

39. Jumper, J. et al. (2021) “Highly accurate protein structure prediction with AlphaFold,” Nature, 596(7873), pp. 583–589. doi: 10.1038/s41586-021-03819-2.

40. Kaplan, C. W. et al. (2005) “Fusobacterium nucleatum apoptosis-inducing outer membrane protein.”

41. Kaplan, C. W. et al. (2009) “The Fusobacterium nucleatum outer membrane protein RadD is an arginine-inhibitable adhesin required for inter-species adherence and the structured architecture of multispecies biofilm.” NIH Public Access.

42. Kaplan, C. W. et al. (2010) “Fusobacterium nucleatum outer membrane proteins Fap2 and RadD induce cell death in human lymphocytes,” Infection and Immunity, 78, pp. 4773–4778.

43. Kidmose, R. T. et al. (2019) “Namdinator – automatic molecular dynamics flexible fitting of structural models into cryo-EM and crystallography experimental maps,” IUCrJ, 6(4), pp. 526–531. doi: 10.1107/S2052252519007619.

44. Kostic, A. D. et al. (2012) “Genomic analysis identifies association of Fusobacterium with colorectal carcinoma.”

45. Kostic, A. D. et al. (2013) “Fusobacterium nucleatum Potentiates Intestinal Tumorigenesis and Modulates the Tumor-Immune Microenvironment.” Cell Press.

46. Lamichhane, S. et al. (2021) “Linking Gut Microbiome and Lipid Metabolism: Moving beyond Associations.” Metabolites.

47. Lattemann, C. T. et al. (2000) “Autodisplay: functional display of active beta-lactamase on the surface of Escherichia coli by the AIDA-I autotransporter,” J Bacteriol, 182(13), pp. 3726–33. doi: 10.1128/JB.182.13.3726-3733.2000.

48. Leo, J. C. et al. (2008) “The Yersinia adhesin YadA binds to a collagenous triple-helical conformation but without sequence specificity,” Protein Engineering, Design and Selection, 21(8), pp. 475–484. doi: 10.1093/protein/gzn025.

49. Levy, M. et al. (2017) “Dysbiosis and the immune system,” Nature Reviews Immunology, 17, pp. 219–232. doi: 10.1038/nri.2017.7.

50. Lima, B. P., Shi, W. and Lux, R. (2017) “Identification and characterization of a novel Fusobacterium nucleatum adhesin involved in physical interaction and biofilm formation with Streptococcus gordonii,” MicrobiologyOpen, 6, p. e00444. doi: 10.1002/mbo3.444.

51. Lin, Y.-X. et al. (2021) “The N-linked glycosylations of TIGIT Asn32 and Asn101 facilitate PVR/TIGIT interaction,” Biochemical and Biophysical Research Communications, 562, pp. 9–14. doi: 10.1016/j.bbrc.2021.05.034.

52. Liu, P. F. et al. (2010) “Vaccination targeting surface FomA of Fusobacterium nucleatum against bacterial co-aggregation: Implication for treatment of periodontal infection and halitosis.”

53. Loftus, M., Hassouneh, S. A.-D. and Yooseph, S. (2021) “Bacterial community structure alterations within the colorectal cancer gut microbiome.” BioMed Central.

54. Lozano, E. et al. (2020) “Nectin-2 Expression on Malignant Plasma Cells Is Associated with Better Response to TIGIT Blockade in Multiple Myeloma,” Clinical Cancer Research, 26(17), pp. 4688–4698. doi: 10.1158/1078-0432.CCR-19-3673.

55. Mastronarde, D. N. (2003) “SerialEM: A Program for Automated Tilt Series Acquisition on Tecnai Microscopes Using Prediction of Specimen Position,” Microscopy and Microanalysis, 9(S02), pp. 1182–1183. doi: 10.1017/S1431927603445911.

56. Melia, C. E. et al. (2021) “Architecture of cell-cell junctions in situ reveals a mechanism for bacterial biofilm inhibition,” Proc Natl Acad Sci U S A, 118(31). doi: 10.1073/pnas.2109940118.

57. Montalban-Arques, A. and Scharl, M. (2019) “Intestinal microbiota and colorectal carcinoma: Implications for pathogenesis, diagnosis, and therapy.”

58. Parhi, L. et al. (2020) “Breast cancer colonization by Fusobacterium nucleatum accelerates tumor growth and metastatic progression,” Nat Commun, 11(1), p. 3259. doi: 10.1038/s41467-020-16967-2.

59. Parhi, L. et al. (2022) “Placental colonization by Fusobacterium nucleatum is mediated by binding of the Fap2 lectin to placentally displayed Gal-GalNAc,” Cell Rep, 38(12), p. 110537. doi: 10.1016/j.celrep.2022.110537.

60. Patil, I. (2021) “Visualizations with statistical details: The ‘ggstatsplot’ approach,” Journal of Open Source Software, 6(61), p. 3167. doi: 10.21105/joss.03167.

61. Paxman, J. J. et al. (2019) “Unique structural features of a bacterial autotransporter adhesin suggest mechanisms for interaction with host macromolecules,” Nature Communications, 10(1), p. 1967. doi: 10.1038/s41467-019-09814-6.

62. Pettersen, E. F. et al. (2004) “UCSF Chimera - A visualization system for exploratory research and analysis,” Journal of Computational Chemistry, 25, pp. 1605–1612. doi: 10.1002/jcc.20084.

63. Pettersen, E. F. et al. (2021) “UCSF ChimeraX: Structure visualization for researchers, educators, and developers,” Protein Sci, 30(1), pp. 70–82. doi: 10.1002/pro.3943.

64. Punjani, A. et al. (2017) “cryoSPARC: algorithms for rapid unsupervised cryo-EM structure determination,” Nat Methods, 14(3), pp. 290–296. doi: 10.1038/nmeth.4169.

65. Rubinstein, M. R. et al. (2013) “Fusobacterium nucleatum Promotes Colorectal Carcinogenesis by Modulating E-Cadherin/β-Catenin Signaling via its FadA Adhesin.” Cell Press.

66. Rubinstein, M. R. et al. (2019) “Fusobacterium nucleatum promotes colorectal cancer by inducing Wnt/beta-catenin modulator Annexin A1,” EMBO Rep, 20(4). doi: 10.15252/embr.201847638.

67. Saha, S. et al. (2023) “The IgV domain of the poliovirus receptor alone is immunosuppressive and binds to its receptors with comparable affinity,” Scientific Reports, 13(1), p. 4609. doi: 10.1038/s41598-023-30999-w.

68. Sanchez-Garcia, R. et al. (2021) “DeepEMhancer: a deep learning solution for cryo-EM volume post-processing,” Commun Biol, 4(1), p. 874. doi: 10.1038/s42003-021-02399-1.

69. Sanders, B. E. et al. (2018) “FusoPortal: an Interactive Repository of Hybrid MinION-Sequenced Fusobacterium Genomes Improves Gene Identification and Characterization.” American Society for Microbiology.

70. Scheres, S.H.W. (2012) “RELION: Implementation of a Bayesian approach to cryo-EM structure determination.” J. Struct. Biol. 180, p. 519–530. doi: 10.1016/j.jsb.2012.09.006.

71. Sender, R., Fuchs, S. and Milo, R. (2016) “Revised Estimates for the Number of Human and Bacteria Cells in the Body,” PLoS Biology, 14.

72. Shhadeh, A. et al. (2021) “CEACAM1 Activation by CbpF-Expressing E. coli,” Front Cell Infect Microbiol, 11, p. 699015. doi: 10.3389/fcimb.2021.699015.

73. Stanietsky, N. et al. (2009) “The interaction of TIGIT with PVR and PVRL2 inhibits human NK cell cytotoxicity.” National Academy of Sciences.

74. Stengel, K. F. et al. (2012) “Structure of TIGIT immunoreceptor bound to poliovirus receptor reveals a cell-cell adhesion and signaling mechanism that requires cis-trans receptor clustering.”

75. Su, Y.-C. et al. (2016) “Haemophilus influenzae P4 Interacts With Extracellular Matrix Proteins Promoting Adhesion and Serum Resistance,” The Journal of Infectious Diseases, 213(2), pp. 314–323. doi: 10.1093/infdis/jiv374.

76. Sung, H. et al. (2021) “Global Cancer Statistics 2020: GLOBOCAN Estimates of Incidence and Mortality Worldwide for 36 Cancers in 185 Countries,” CA Cancer J Clin, 71(3), pp. 209–249. doi: 10.3322/caac.21660.

77. Tegunov, D. and Cramer, P. (2019) “Real-time cryo-electron microscopy data preprocessing with Warp,” Nat Methods, 16(11), pp. 1146–1152. doi: 10.1038/s41592-019-0580-y.

78. Tierney, B. T. et al. (2019) “The Landscape of Genetic Content in the Gut and Oral Human Microbiome,” Cell Host and Microbe, 26, pp. 283–295.

79. Valguarnera, E. and Wardenburg, J. B. (2020) “Good Gone Bad: One Toxin Away From Disease for Bacteroides fragilis.” Elsevier Ltd.

80. Vinayagam, D. et al. (2018) “Electron cryo-microscopy structure of the canonical TRPC4 ion channel,” Elife, 7. doi: 10.7554/eLife.36615.

81. Wilson, M. R. et al. (2019) “The human gut bacterial genotoxin colibactin alkylates DNA,” Science, 363, p. eaar7785. doi: 10.1126/science.aar7785.

82. Wirbel, J. et al. (2019) “Meta-analysis of fecal metagenomes reveals global microbial signatures that are specific for colorectal cancer.” Nature Publishing Group.

83. Wu, L. et al. (2021) “Altered Gut Microbial Metabolites in Amnestic Mild Cognitive Impairment and Alzheimer’s Disease: Signals in Host–Microbe Interplay.” Nutrients.

84. Xu, M. et al. (2007) “FadA from Fusobacterium nucleatum utilizes both secreted and nonsecreted forms for functional oligomerization for attachment and invasion of host cells.” J Biol Chem.

85. Yang, G. Y. and Shamsuddin, A. M. (1996) Gal-GalNAc: A biomarker of colon carcinogenesis, Histology and Histopathology.

86. Yu, X. et al. (2009) “The surface protein TIGIT suppresses T cell activation by promoting the generation of mature immunoregulatory dendritic cells.” Nature Publishing Group.

87. Yuan, B. et al. (2021) “Fecal Bacteria as Non-Invasive Biomarkers for Colorectal Adenocarcinoma,” Front Oncol, 11, p. 664321. doi: 10.3389/fonc.2021.664321.

88. Zhang, Q. et al. (2018) “Blockade of the checkpoint receptor TIGIT prevents NK cell exhaustion and elicits potent anti-tumor immunity,” Nat Immunol, 19(7), pp. 723–732. doi: 10.1038/s41590-018-0132-0.

89. Zondlo, N. J. (2013) “Aromatic-proline interactions: electronically tunable CH/π interactions,” Accounts of Chemical Research, 46(4), pp. 1039–1049. doi: 10.1021/ar300087y.

90. van Zundert, G. C. P. et al. (2016) “The HADDOCK2.2 Web Server: User-Friendly Integrative Modeling of Biomolecular Complexes,” Journal of Molecular Biology, 428(4), pp. 720–725. doi: 10.1016/j.jmb.2015.09.014.

