## Supplementary Information for "Structural basis of *Fusobacterium nucleatum* adhesin Fap2 interaction with receptors on cancer and immune cells"

1: Leibniz Forschungsinstitut für Molekulare Pharmakologie (FMP), Robert-Rössle-Straße 10, 13125 Berlin, Germany

2: Present address: University of Potsdam, Faculty of Science, Karl-Liebknecht-Straße 24-25, 14476 Potsdam OT Golm, Germany

3: Core Facility for Cryo Electron Microscopy of the Charité - Universitätsmedizin Berlin at the Max Delbrück Center, Robert-Rössle-Straße 10, 13125 Berlin, Germany

4: Max Delbrück Center for Molecular Medicine, Technology Platform Cryo-EM, Robert-Rössle-Straße 10, 13125 Berlin, Germany

5: Institute of Microbiology, Infectious Diseases and Immunology, Biofilmcenter, Charité - Universitätsmedizin Berlin, Hindenburgdamm 30, 12203 Berlin, Germany

**Supplementary figures.**

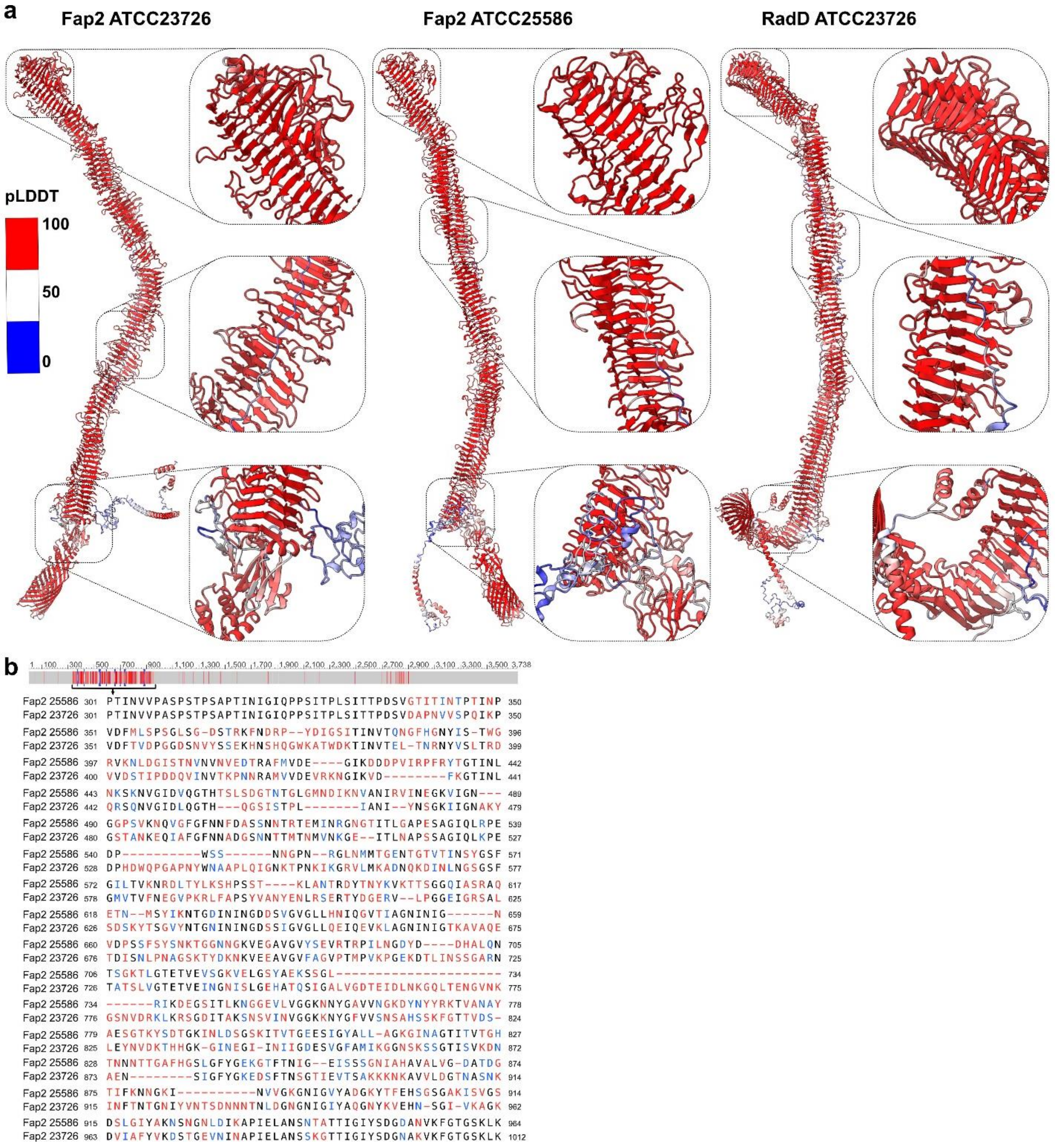


Supplementary Figure 1: AlphaFold2 prediction and sequence alignment of Fap2 and RadD. a) Structure model prediction of adhesins Fap2 from Fn strains ATCC23726 and ATCC25586 and RadD from strain ATCC23726. Models are colored according to prediction reliability (pLDDT). Sections of the matchstick head, the β-helix with the unstructured N-terminus, and the C-terminal end of the β-helix are magnified. b) Sequence alignment of the N-terminal part of the β-helix including part of the unstructured region (residues 301-364) of both Fap2, colored by conservation.

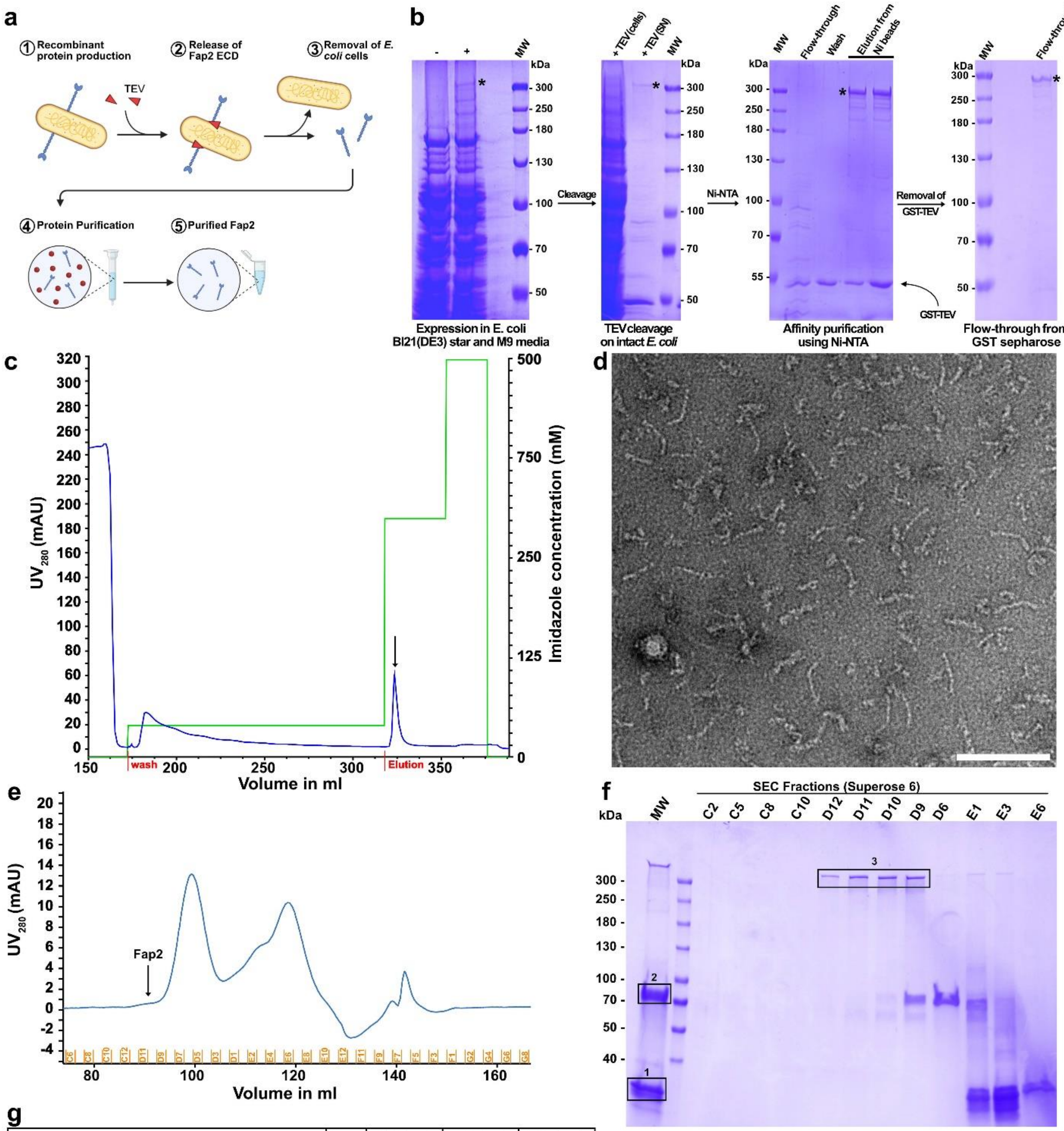


g

| Protein names | Gene names | LFQ intensity 1 | LFQ intensity 2 | LFQ intensity 3 |
|---|---|---|---|---|
| Bifunctional polymyxin resistance protein ArnA | arnA | 188760000 | 8802600000 | 110060000 |
| FKBP-type peptidyl-prolyl cis-trans isomerase SlyD | slyD | 9124600000 | 41709000 | 0 |
| Glutamine--fructose-6-phosphate aminotransferase [isomerizing] | glmS | 14908000 | 2808200000 | 21430000 |
| Ubiquinone/menaquinone biosynthesis C-methyltransferase UbiE | ubiE | 0 | 0 | 0 |
| **Fap2-ECD** | **fap2** | **11986000** | **4618700** | **2017500000** |
| **Tev** | **tev** | **7011600000** | **76486000** | **4315400** |
| Glycerol-3-phosphate regulon repressor | glpR | 135400000 | 311850 | 0 |
| cAMP-activated global transcriptional regulator CRP | crp | 0 | 0 | 0 |
| Ribosomal small subunit pseudouridine synthase A | rsuA | 114570000 | 204570 | 0 |
| Chaperone protein DnaK | dnaK | 0 | 35483000 | 177040 |
| L-fucose operon activator | fucR | 0 | 0 | 0 |
| Uncharacterized HTH-type transcriptional regulator YihW | yihW | 0 | 0 | 0 |
| NAD kinase | nadK | 0 | 0 | 0 |
| 60 kDa chaperonin | groL | 383660 | 13445000 | 4467600 |
| 4-hydroxy-tetrahydrodipicolinate reductase | dapB | 0 | 0 | 0 |

Supplementary Figure 2: Purification strategy of Fap2-ECD. a) Schematics of the work flow of Fap2-ECD expression in fusion with the AIDA autotransporter in the *E. coli* outer membrane, followed by release through TEV cleavage and purification. b) SDS-PAGEs showing the expression, release by TEV, and the first two purification steps. Fap2-ECD is indicated by *. c)

Chromatogram showing the IMAC purification of Fap2-ECD after TEV cleavage. The arrow indicates the main peak of the elution, which has been visualized in d). d) Negative stain EM micrograph of Fap2-ECD after IMAC as indicated in c). Scale bar: 100 nm. e) Chromatogram showing size exclusion chromatography (SEC) of Fap2-ECD on Superose 6. f) SDS-PAGE of SEC run in e). Proteins have been concentrated 10x through TCA precipitation before loading on the gel. The three indicated bands were analyzed by peptide fingerprint in g). g) Peptide fingerprint mass spectrometry analysis of the three protein bands indicated in f). Fap2 is the main constituent of sample 3, whereas TEV is the main protein in sample 1.

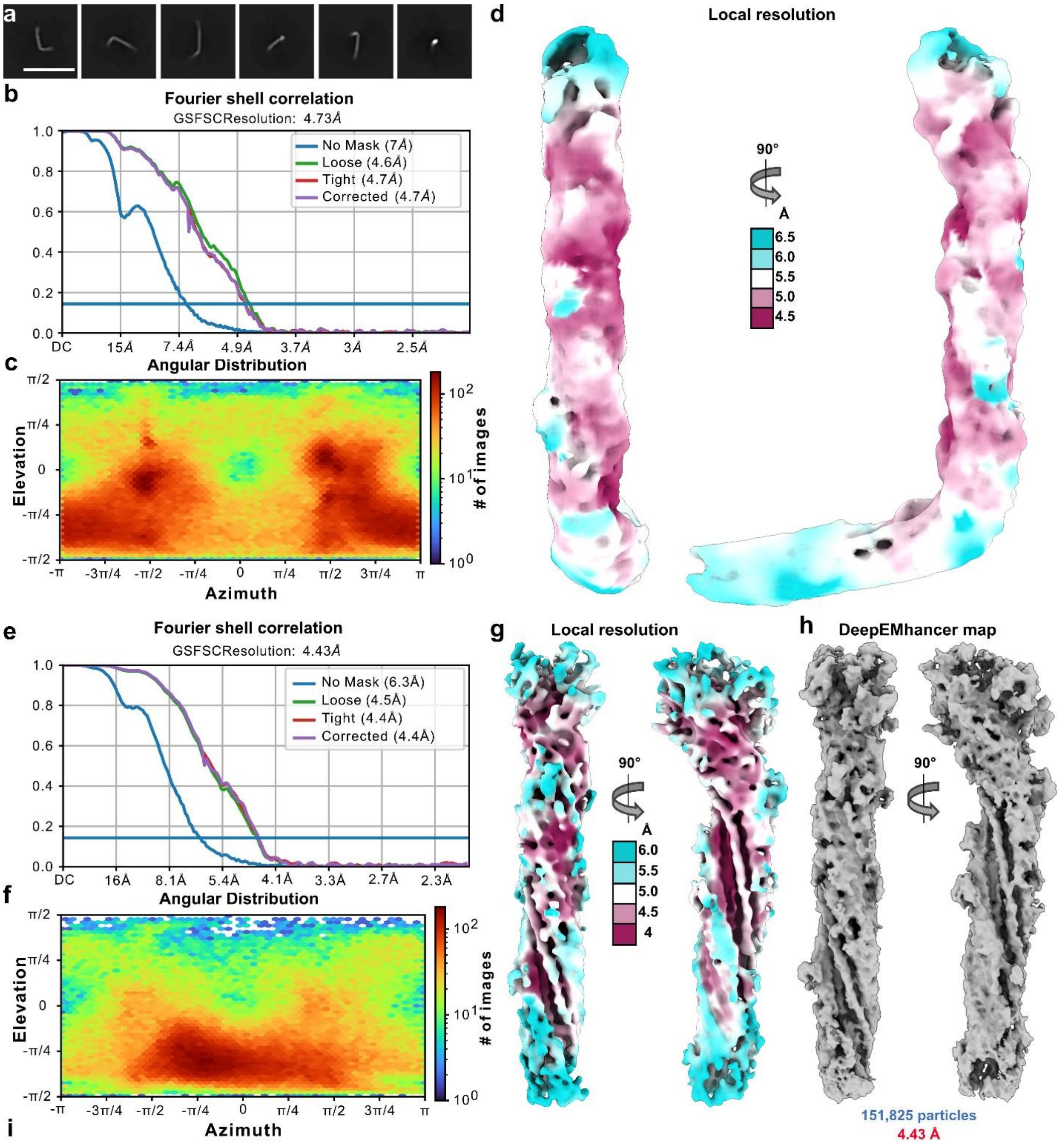


i

```
Query                            1    NKSWGLVDLRDLREPTNEVEILAHISPKEVTKQVISLN--ITEPTVQTLEAPDVNPQVNEPLPAPVIELPEVETVTI---NPLSINAPSAPTAPSAPT 92
F. nucleatum ATCC23726           195  NKSWGLVDLRDLREPTNEVEILAHISPKEVTKQVISLN--ITEPTVQTLEAPDVNPQVNEPLPAPVIELPEVETVTI---NPLSINAPSAPTAPSAPT 286
F. vincentii                     203  NRSWGLVDLKDLKEPTNEVEILARISPKEVTKQAISLN--ITEPTVQTLEAPTVNPQVNEPLPAPVIELPEVETVTI---NPLTINAPSAPTAPSAPT 294
F. nucleatum ATCC25586           195  NKSWGLVDLRDLREPTNEVEILAHISPKEVTKQAISLN--ITEPTVQTLEAPDVNPQVNEPLPAPVIELPEVETVTI---NPLSINAPGAPTAPSAPT 286
F. animalis                      203  NRNWGLVDLKDLREPTNEVEILARISPKEVTKQAISLN--ITEPTVQTLEAPAVNPQVNEPLPAPVIELPEVETVTI---NPLTINAPSAPTAPNAPS 294
F. polymorphum                   195  NRSWGLVDLKDLKEPTNEVEILANISPKEVTKQAISLN--ITEPTVQTLEAPDVNPQVNEPLPAPVIDLPEVETVTI---NPLTINAPSAPSAPTAPN 286
Pseudoleptotrichia goodfellowii  200  NRSWGLVDLRNLKEPTSEVEILARISPKEVTKQAITLN--ITQPAINELEAPIVNPQVNKPLDAPVITLPEVEKVEI---SALNITAPSAPTAPTAPG 291
F. periodonticum                 197  NRTWGLVDLLNLKEPTNEVEILARISPKEVSKQALALN--ITAPVVQTLEAPVVKPEPNKPIEAPKINLPSIKKVTI---EALSINPPSAPGVIGVPS 288
Leptotrichia wadei               168   QSYGLASLLHVQEPEVEIQIMANVRPKSVSKEEIAINPKIDMP--REVVRPNINLKVTEPITAPNITFPNVKPINIDIKNPV---APSEPKAIGAPN 256

 Query                           93   ---INI---------AITAPSAPVP---------PTINVVPASPSTPSAPTINIGIQPPSITPLSITTPDSVDA--PNVVSPQIKPVDFTVDPGGDSN 168
F. nucleatum ATCC23726           287  ---INI---------AITAPSAPVP---------PTINVVPASPSTPSAPTINIGIQPPSITPLSITTPDSVDA--PNVVSPQIKPVDFTVDPGGDSN 362
F. vincentii                     295  ---INI---------GITAPSAPIP---------PSINVVPASPSTPSAPSINIGVQPPSITPLSITTPDSVDA--PNVVSPQIKPVDFTVDPGGDSN 370
F. nucleatum ATCC25586           287  ---INI---------GITAPSAPSA---------PTINVVPTSPSTPSAPSINIGIQPPRITPLSITTPDSVGT--ITINTPTINPVDFMLSPSG    359
F. animalis                      295  ---ISI---------GINAPSAPVS---------PTISVVPTSPSAPSAPTINIGIQPPSITALSISTPSTVVP--PEVVSPNITPVDFVIDPAGNS  369
F. polymorphum                   287  ---ISI---------GITAPTAPTAPSAPIAPEVPTITVNPSSPETVSAPTININVTIPKISTLDITTPGSVGT--ISVIAPTVTPVDF          362
Pseudoleptotrichia goodfellowii  292  APVINISITPPNAPNALTAPSAPSV---------PNISVTPSAPSVLTAPNININVTAPSINALTIATPPTVVA--PQVVSPQIKPVDFVIDPSGDA  378
F. periodonticum                 289  ---IGI---------TIATPSKPVP---------PTVSVSVNEPSAPGAPTINIPVTPPKIVALNIETPPAVSVTPPTV--SAIKPVAFSVDPTIDSN 364
Leptotrichia wadei               257  ---INI---------TLEAP----Q---------VTLGITP-----PKTPNINISVTPPDITPLTIAKPSEVDL--PTVTPPTINPVDFSIDPTGDS  324
```

Supplementary Figure 3: Cryo-EM and SPA of Fap2-ECD. a) Representative 2D class averages, showing Fap2 in side view and front view. Scale bar: 50 nm. b-d) Fourier shell correlation (b), angular distribution heatmap (c), and density map colored by local resolution (d) of the final cryo-EM density of the full Fap2-ECD. e-h) Fourier shell correlation (e), angular distribution heatmap (f), density map colored by local resolution (g), and final density map sharpened with

DeepEMhancer of the locally refined longer branch of Fap2-ECD including the matchstick region (h). i) Sequence alignment of the N-terminus of Fap2 (residues 195 - 362) from Fn ATCC23726 with related autotransporter sequences from other *Fusobacteriota*.

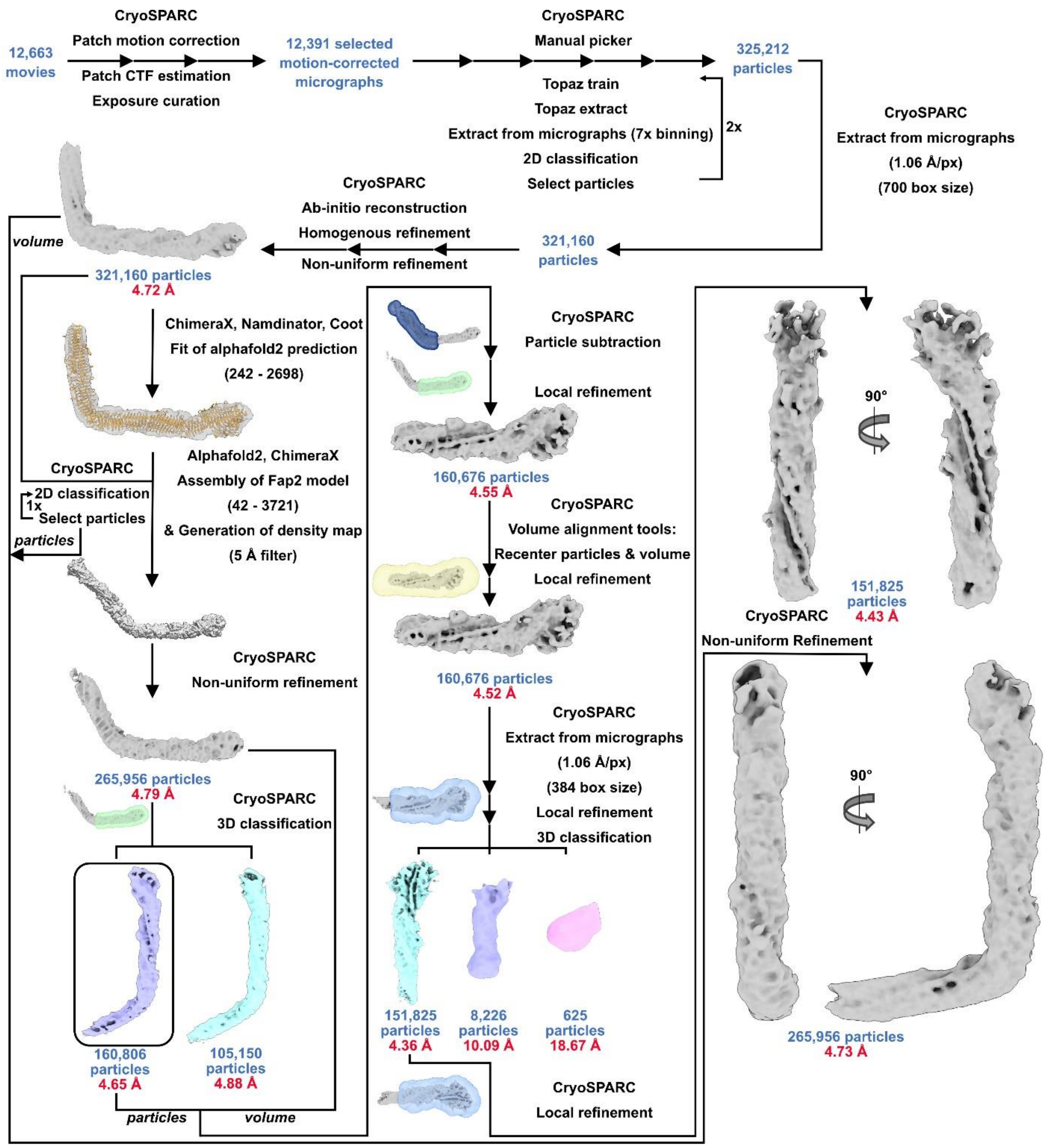


Supplementary Figure 4: SPA data processing scheme of the Fap2-ECD, as shown in SI Fig. 3. Masks for signal subtraction or local refinements are shown transparent.

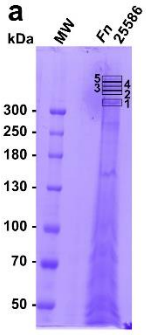


b

| Protein names | Gene names | LFQ intensity 1 | LFQ intensity 2 | LFQ intensity 3 | LFQ intensity 4 | LFQ intensity 5 |
|---|---|---|---|---|---|---|
| Haloalkane dehalogenase | dhaA | 0 | 0 | 6663100000 | 28724000 | 0 |
| Methylated-DNA-protein-cysteine methyltransferase | MGMT | 0 | 0 | 4622000000 | 0 | 0 |
| **RadD** | **RadD** | **12523000000** | **271020000** | **45087000** | **191520000** | **93909000** |
| Trigger factor | tig | 270220000 | 227600000 | 75328000 | 720420000 | 363890000 |
| Dermcidin;Survival-promoting peptide;DCD-1 | DCD | 0 | 0 | 0 | 0 | 4074100 |
| Ubiquitin related | Ubiquitin related | 0 | 30197000 | 8575000 | 71303000 | 37108000 |
| **Fap2** | **Fap2** | **293800000** | **205270000** | **51167000** | **824620000** | **375500000** |
| Glutathione-S-Transferase | GST 26 | 0 | 9299400 | 0 | 0 | 0 |
| Cystatin-A;Cystatin-A, N-terminally processed | CSTA | 0 | 0 | 6059600 | 0 | 0 |
| Transthyretin | TTR | 0 | 0 | 41707000 | 0 | 0 |
| Ubiquitin related modifying proteins | Ubiquitin related | 0 | 0 | 0 | 0 | 2436800 |
| Calmodulin-like protein 5 | CALML5 | 0 | 8275700 | 5953900 | 0 | 5531100 |
| Chaperone protein DnaK | dnaK | 4666300 | 3624900 | 158540000 | 12469000 | 5716800 |
| Basic salivary proline-rich protein 1;Proline-rich peptide II-2;Basic peptide IB-6;Peptide P | PRB1 | 0 | 22626000 | 2194600 | 1190400 | 0 |
| BPI fold-containing family A member 1 | BPIFA1 | 0 | 7847200 | 0 | 0 | 0 |

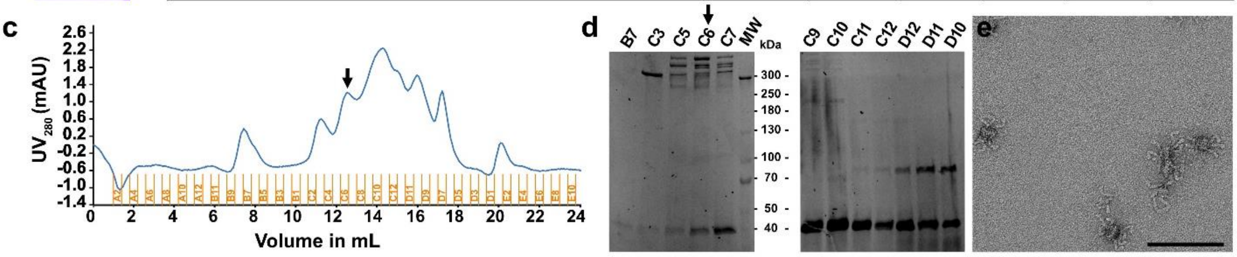


Supplementary Figure 5: Extraction and purification of native Fap2 from Fn ATCC25586. a) SDS-PAGE showing total protein content of Fn ATCC25586. The five indicated bands were analyzed by peptide fingerprint in (b). b) Peptide fingerprint mass spectrometry analysis of the five protein bands indicated in (a). Fap2 is the main protein in samples 4 and 5, whereas RadD is the main protein in sample 1. Both proteins are nearly equally present in samples 3 and 4. c,d) Chromatogram (c) and SDS-PAGE (d) showing SEC of extracted OM proteins of Fn on Superose 6. e) Negative stain EM micrograph of SEC fraction C6 (indicated by arrow in d). Note the rod-shaped molecules that cluster together. Scale bar: 100 nm.

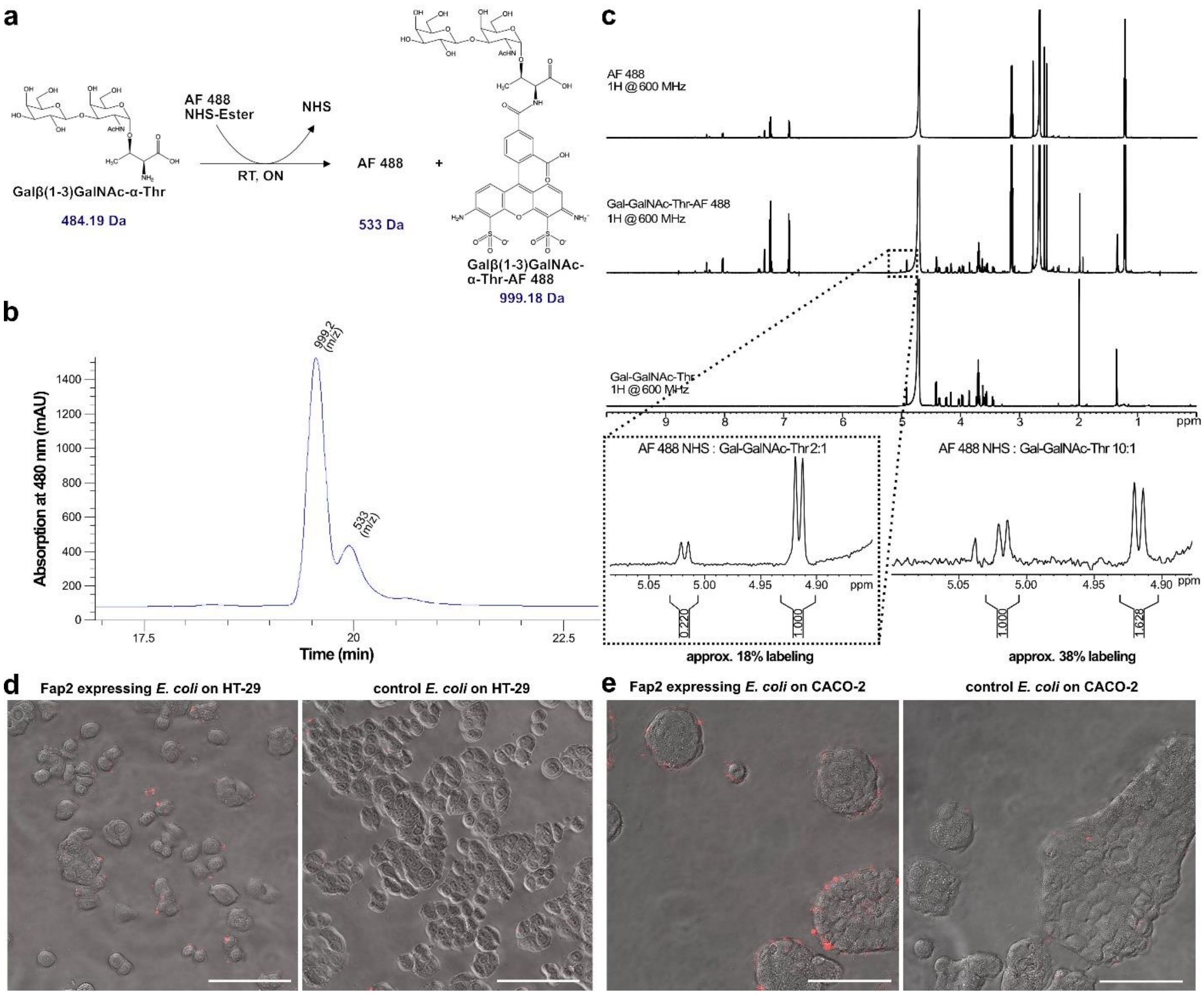


Supplementary Figure 6: Preparation of fluorescently labeled Gal-GalNAc-Thr and CRC cell binding of Fap2-expressing *E. coli*. a) Scheme of the labeling reaction, including the expected masses of the reactants and products. b) Chromatogram showing LC-MS of Gal-GalNAc-Thr after labeling with AF 488 NHS-Ester (Lumiprobe) The mass of the main peak is in agreement with the expected product. c) 1H-NMR spectrum of fluorescently labeled Gal-GalNAc-Thr. The inset shows two characteristic peaks for the educt and product at two ratios of the reactants. d,e) Micrographs that show binding to HT-29 (d) or Caco-2 (e) colon cancer cells of Fap2-expressing *E. coli*. Red channel: *E. coli* labeled with CellBrite Fix 555, gray: bright field image. Scale bars: 50 µm.

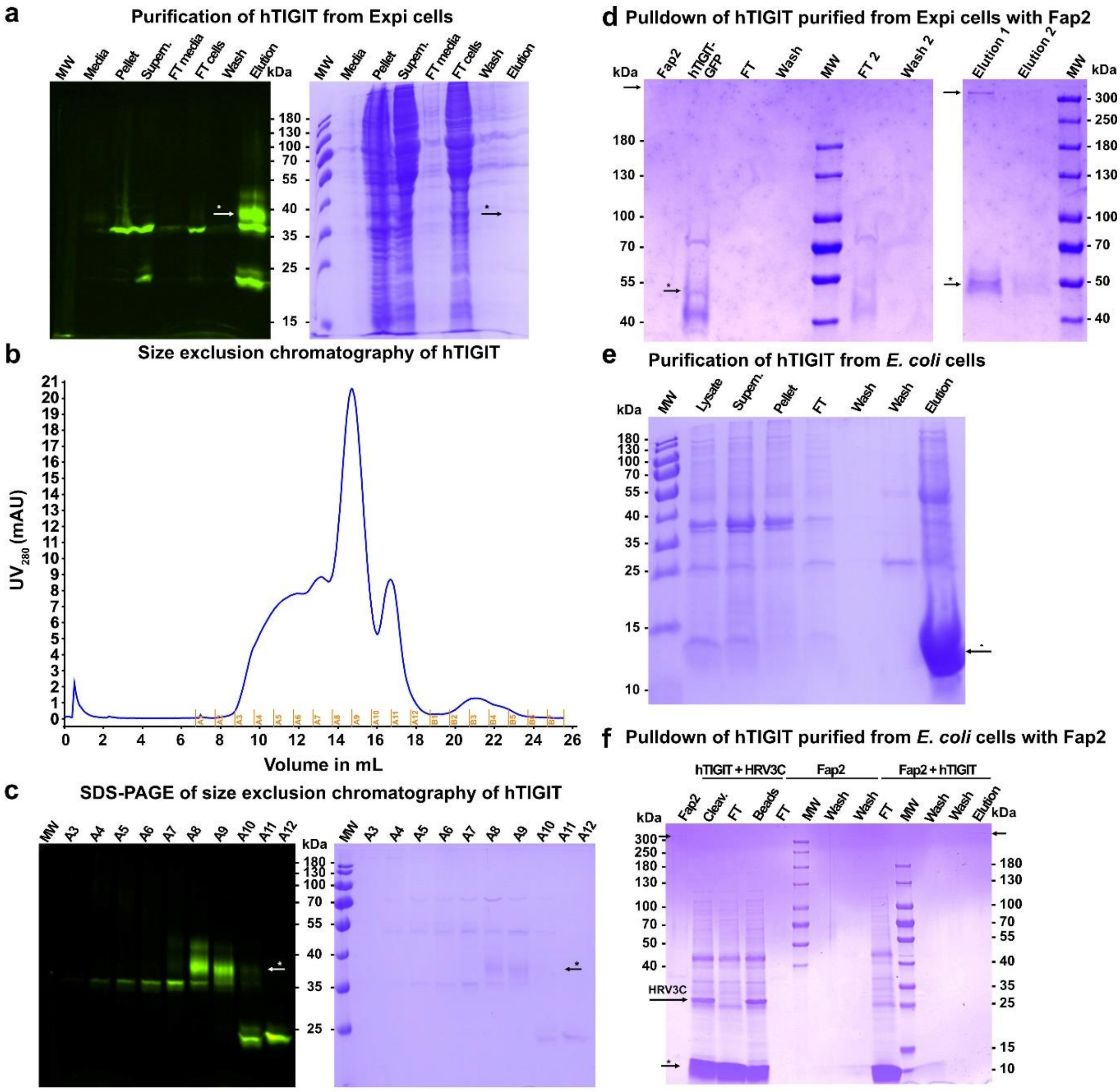


Supplementary Figure 7: Purification of hTIGIT-ECD and pulldown with Fap2. a) SDS-PAGE showing purification of hTIGIT-ECD as fusion protein with GFP (hTIGIT-ECD-GFP) from Expi cells. Left: in-gel fluorescence of GFP, right: coomassie staining. hTIGIT-ECD is indicated. b) Chromatogram showing SEC of hTIGIT-ECD-GFP as last purification step. c) SDS-PAGE of the SEC in (b). The position of hTIGIT-ECD-GFP in fractions A8/A9 is indicated. d) SDS-PAGE showing pulldown of hTIGIT-ECD-GFP (a-c) with Fap2-ECD. The positions of Fap2-ECD and hTIGIT-ECD-GFP are indicated (bands above 300 kDa and at 50 kDa, respectively). e) Purification of hTIGIT-ECD that have been insolubly expressed in *E. coli* and refolded, as described in (Stengel et al., 2012). f) SDS-PAGE showing pulldown of hTIGIT-ECD purified from *E. coli* (e) with Fap2. The positions of hTIGIT-ECD (arrow with *) and Fap2-ECD (arrow) are indicated. No visible band of hTIGIT appears in the pulldown elution fraction.

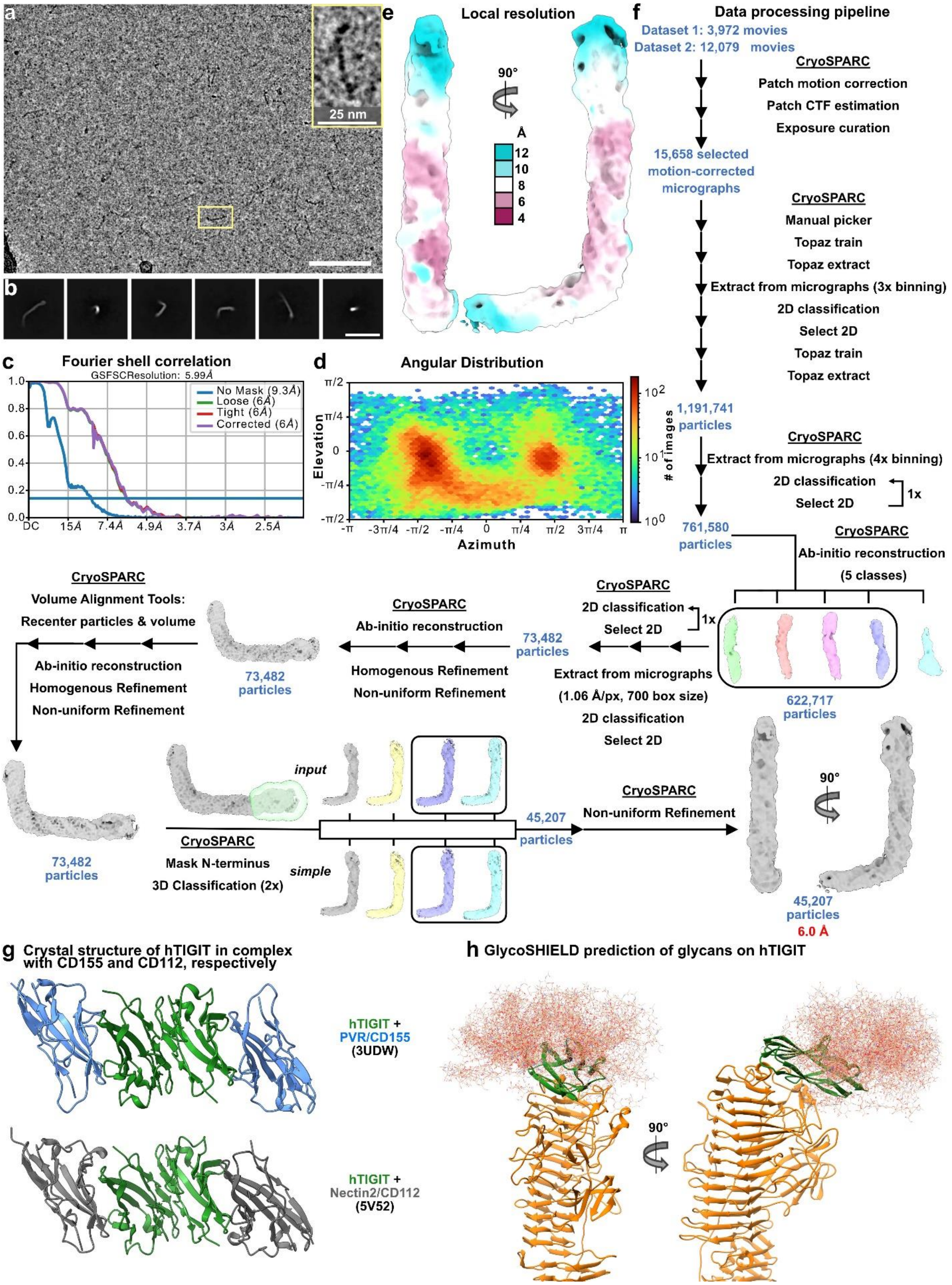


Supplementary Figure 8: Cryo-EM and SPA of Fap2-ECD/hTIGIT-ECD complex. a) Representative cryo-EM micrograph recorded at 300 kV and -2.0 µm defocus. Scale bar: 100 nm. Inset: one molecule in magnified view. b) Representative 2D class averages, showing the complex in side view and front view. Scale bar: 50 nm. c,d,e) Fourier shell correlation (c), angular distribution heatmap (d), and density map colored by local resolution (e) of the final cryo-EM density of the complex. f) SPA data processing scheme of the complex. g) Crystal structures of heterotetramers of hTIGIT-ECD with PVR (3UDW) and Nectin2 (5V52). h) In silico glycosylation of the hTIGIT-ECD model in complex with Fap2. Glycan poses were obtained using GlycoSHIELD (GlycoSHIELD: a versatile pipeline to assess glycan impact on protein structures | bioRxiv, no date). 40 glycan poses (complex biantennal glycan) are shown. According to this model, glycans are not directly involved in the Fap2/hTIGIT interface.

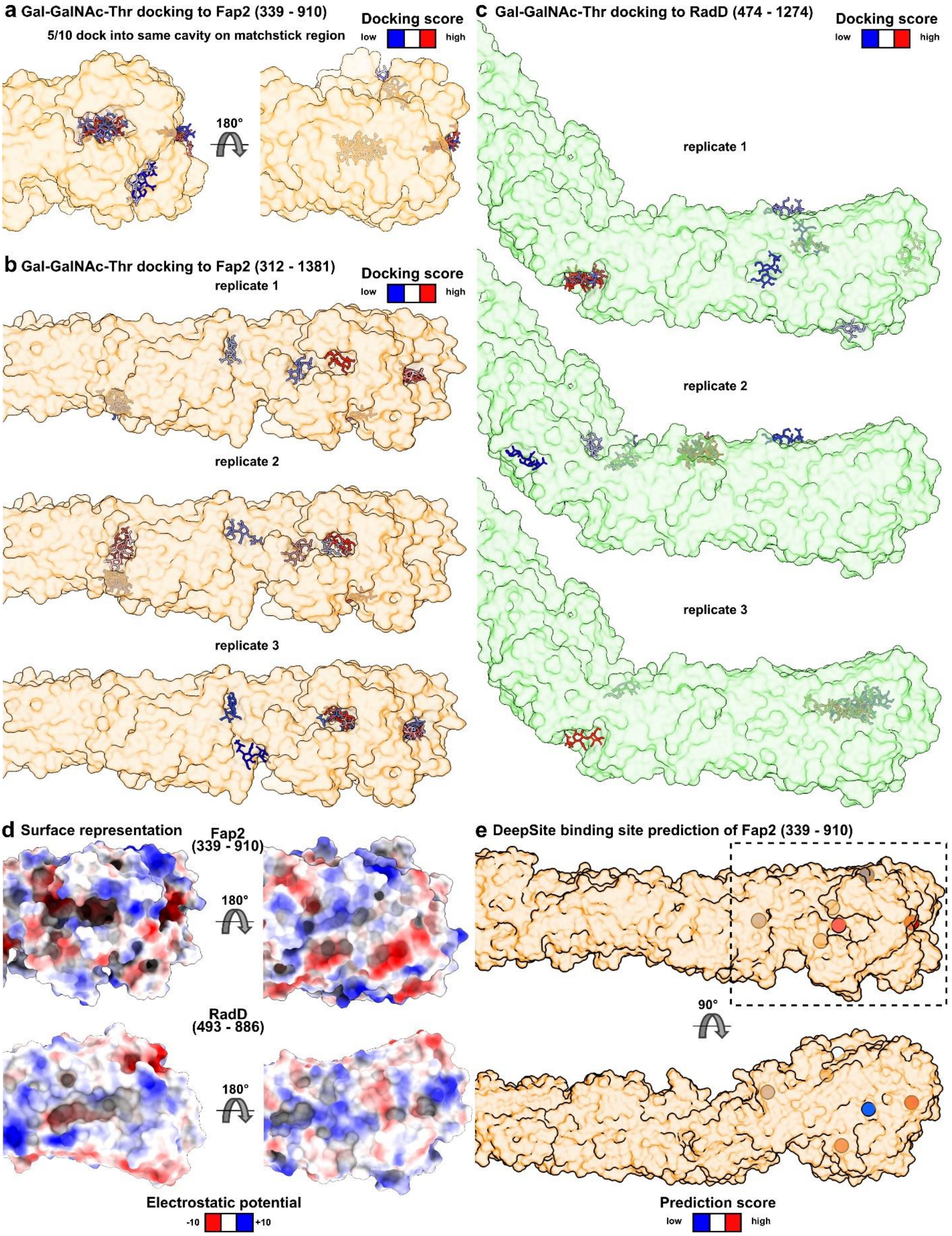


Supplementary Figure 9: Docking of Gal-GalNAc-Thr to Fap2 and RadD. a) Docking of Gal-GalNAc-Thr to the matchstick region of Fap2. 5 out of 10 poses are docked into the proposed binding grove. b,c) Three docking replicates of Gal-GalNAc-Thr to the tips of the β-helices of Fap2 (b) and RadD (c). For Fap2, the docking poses with the highest scores are always docked in the proposed binding grove. For RadD, the docking poses are randomly distributed. The Autodock Vina plugin in UCSF Chimera was used for dockings in (a-c). d) Surface representations of the

Fap2 and RadD matchstick regions, colored by electrostatic potential. e) Binding site prediction on the Fap2 matchstick region by DeepSite (Jiménez et al., 2017). The two out of five highest prediction scores (red dots) revealed the proposed binding sites for Gal-GalNAc and TIGIT.

**Supplementary Tables.**

Supplementary Table 1: Cryo-EM data collection and refinement statistics.

| | Fap2 (Res. 42-3271) | Fap2 (Res. 42-3271) + hTIGIT (1-141)-GFP |
|---|---|---|
| **Data collection and processing** | | |
| Magnification | 81,000 | 81,000 |
| Voltage (kV) | 300 | 300 |
| Camera | Bioquantum K3 | Bioquantum K3 |
| Electron exposure (e–/Å$^2$) | 59.8 | 44.6 |
| Defocus range (μm) | -1.5 – 2.6 | -1.0 – 2.5 |
| Pixel size (Å) | 0.53 | 0.53 |
| Micrographs used | 12,391 | 15,658 |
| Total extracted particle images | 375,308 | 1,191,741 |
| Refined particle images | 321,160 | 761,580 |
| Final particle images | 151,825 (head)<br>265,956 (full) | 45,207 |
| Map resolution, head (Å) | 4.4 (head); 4.7 (full) | 6.0 |
| FSC threshold | 0.143 | 0.143 |
| Map resolution range (Å) | 3.7 – 7.2 (head)<br>4.1 – 13.3 (full) | 4.6 – 14.1 |

Supplementary Table 2: SPR statistics of hTIGIT-Fc binding to Fap2-ECD

**Parameters**

| Sample | Conc | Rmax | SE(Rmax) | ka | SE(ka) | kd | SE(kd) | RI | SE(RI) | ka (1/Ms) | kd (1/s) | Rmax (RU) | RI (RU) | KA (1/M) |
|---|---|---|---|---|---|---|---|---|---|---|---|---|---|---|
| | | 36.2 | 1.95 | 5.29E+04 | 3.12E+03 | 0.0308 | 1.10E-03 | | | 5.29e4 | 0.0308 | 36.2 | | 1.72e6 5.8[illegible] |
| TIGIT-Fc + 52,5 nM Fap-2 | 52.5n | | | | | | | -1.38 | 0.0777 | | | | -1.38 | |
| TIGIT-Fc + 148 nM Fap-2 | 148n | | | | | | | 0.243 | 0.132 | | | | 0.243 | |
| TIGIT-Fc + 105 nM Fap-2 | 105n | | | | | | | -0.684 | 0.109 | | | | -0.684 | |
| TIGIT-Fc + 368 nM Fap-2 | 368n | | | | | | | 5.36 | 0.245 | | | | 5.36 | |

| Sample | KD (M) | Req (RU) | kobs (1/s) | Chi2 |
|---|---|---|---|---|
| | e-7 | | | 0.477 |
| TIGIT-Fc + 52,5 nM Fap-2 | | 2.99 | 0.0336 | |
| TIGIT-Fc + 148 nM Fap-2 | | 7.33 | 0.0387 | |
| TIGIT-Fc + 105 nM Fap-2 | | 5.52 | 0.0364 | |
| TIGIT-Fc + 368 nM Fap-2 | | 14 | 0.0503 | |

**Parameters**

| Sample | Inject Start | Fit Start | Fit End | Inject End | Fit Start | Fit End |
|---|---|---|---|---|---|---|
| TIGIT-Fc + 52,5 nM Fap-2 | 141 | 141 | 285 | 291 | 291 | 314 |
| TIGIT-Fc + 148 nM Fap-2 | 139 | 141 | 285 | 288 | 291 | 314 |
| TIGIT-Fc + 105 nM Fap-2 | 139 | 141 | 285 | 288 | 291 | 314 |
| TIGIT-Fc + 368 nM Fap-2 | 139 | 141 | 285 | 288 | 291 | 314 |

Supplementary Table 3: Molecular docking of hTIGIT-ECD to Fap2-ECD with HADDOCK2.2

| Parameters | Values |
|---|---|
| HADDOCK score | 21.2 +/- 11.4 |
| Cluster size | 8 |
| RMSD from the overall lowest-energy structure | 0.6 +/- 0.4 |
| Van der Waals energy | -106.4 +/- 3.5 |
| Electrostatic energy | -268.2 +/- 14.2 |
| Desolvation energy | -4.1 +/- 1.7 |
| Restraints violation energy | 1853.5 +/- 112.12 |
| Buried Surface Area | 2957.7 +/- 61.9 |
| Z-Score | -1.9 |

Supplementary Table 4: Occupancy of H-bonds during MD simulation of Fap2-ECD/hTIGIT-ECD complex.

| donor | acceptor | rep 1 | rep 2 | rep 3 | rep 4 | rep 5 | rep 6 | rep 7 | rep 8 | rep 9 | rep 10 | occupancy [%] | standard deviation %] |
|---|---|---|---|---|---|---|---|---|---|---|---|---|---|
| VAL32-Main-N | LYS368-Main-O | 0,97 | 0,99 | 0,99 | 0,98 | 0,99 | 0,98 | 0,98 | 0,79 | 0,94 | 0,98 | 0,96 | 0,06 |
| SER371-Side-OG | ALA30-Main-O | 0,65 | 1,00 | 0,73 | 1,00 | 0,99 | 0,56 | 0,74 | 0,80 | 0,76 | 0,89 | 0,81 | 0,14 |
| TRP540-Side-NE1 | PRO65-Main-O | 0,93 | 0,90 | 0,00 | 0,03 | 0,10 | 0,65 | 0,96 | 0,79 | 0,15 | 0,92 | 0,54 | 0,40 |
| ARG427-Side-NH2 | GLN22-Side-OE1 | 0,33 | 0,49 | 0,20 | 0,50 | 0,39 | 0,31 | 0,09 | 0,38 | 0,06 | 0,14 | 0,29 | 0,15 |
| ARG427-Side-NH1 | HIS24-Side-NE2 | 0,17 | 0,48 | 0,17 | 0,10 | 0,56 | 0,03 | 0,02 | 0,66 | 0,41 | 0,18 | 0,28 | 0,22 |
| TRP53-Main-N | ASN541-Side-OD1 | 0,56 | 0,46 | 0,01 | 0,04 | 0,01 | 0,37 | 0,20 | 0,17 | 0,06 | 0,49 | 0,24 | 0,20 |
| ASN541-Side-ND2 | TRP53-Main-O | 0,22 | 0,19 | 0,00 | 0,01 | 0,01 | 0,31 | 0,26 | 0,18 | 0,06 | 0,40 | 0,17 | 0,13 |
| LEU25-Main-N | HIS372-Side-ND1 | 0,02 | 0,04 | 0,05 | 0,87 | 0,16 | 0,00 | 0,13 | 0,01 | 0,00 | 0,12 | 0,14 | 0,25 |

Supplementary Table 5: Occupancy of H-bonds during MD simulation of Fap2-ECD/Gal-GalNAc-Thr complex.

| donor | acceptor | rep 1 | rep 2 | rep 4 | rep 5 | rep 7 | rep 9 | occupancy [%] | standard deviation [%] |
|---|---|---|---|---|---|---|---|---|---|
| PHE818-Main-N | LIG911-Side-O1 | 80,65 | 99,21 | 0,56 | 98,65 | 2,17 | 84,63 | 60,98 | 46,77 |
| LIG911-Side-O5 | GLU619-Side-OE1 | 21,26 | 82,74 | 0,52 | 58,96 | 51,18 | 56,77 | 45,24 | 29,45 |
| LIG911-Side-O6 | GLU619-Side-OE1 | 23,65 | 43,00 | 10,26 | 71,00 | 67,87 | 17,21 | 38,83 | 26,11 |
| LIG911-Side-O5 | GLU619-Side-OE2 | 14,59 | 17,85 | 11,73 | 46,99 | 66,95 | 60,84 | 36,49 | 24,78 |
| LIG911-Side-O6 | GLU619-Side-OE2 | 13,93 | 23,95 | 13,14 | 61,74 | 59,83 | 13,33 | 30,99 | 23,45 |
| LIG911-Side-N2 | GLU713-Side-OE2 | 75,41 | 15,29 | 0,00 | 68,82 | 0,00 | 0,00 | 26,59 | 35,82 |
| LIG911-Side-N2 | GLU713-Side-CD | 48,12 | 27,42 | 0,00 | 73,54 | 0,00 | 0,00 | 24,85 | 30,89 |
| LYS710-Side-NZ | LIG911-Side-O13 | 4,60 | 8,55 | 11,96 | 3,57 | 10,61 | 87,80 | 21,18 | 32,80 |
| LIG911-Side-N2 | GLU713-Side-OE1 | 16,31 | 43,34 | 0,00 | 59,34 | 0,00 | 0,01 | 19,83 | 25,71 |
| LIG911-Side-O6 | GLU619-Side-CD | 10,39 | 29,65 | 12,98 | 57,66 | 0,00 | 0,00 | 18,45 | 22,08 |
| LEU615-Main-N | LIG911-Side-O5 | 6,51 | 28,68 | 0,15 | 68,31 | 1,34 | 0,04 | 17,51 | 27,18 |
| LIG911-Side-O5 | GLU619-Side-CD | 15,62 | 42,90 | 2,32 | 44,14 | 0,00 | 0,00 | 17,50 | 20,98 |
| LIG911-Side-O4 | ARG613-Main-O | 9,04 | 34,80 | 0,00 | 44,57 | 1,83 | 0,03 | 15,05 | 19,62 |
| LYS710-Side-NZ | LIG911-Side-O12 | 4,39 | 29,51 | 13,75 | 1,51 | 11,20 | 5,53 | 10,98 | 10,14 |
| ARG622-Side-NH2 | LIG911-Side-O6 | 2,84 | 0,64 | 0,11 | 5,86 | 8,26 | 44,04 | 10,29 | 16,82 |

**Supplementary Movie legends.**

Supplementary Movie 1: Structure of the Fap2-ECD. The movie shows a fly-through of longitudinal sections of the density map, highlighting the triangular shaped groove that is complemented by the unstructured N-terminus.

Supplementary Movie 2: Representative MD simulation of the Fap2-ECD/TIGIT-ECD complex. The movie shows a simulation over 50 nsec of the best docking result from HADDOCK. Fap2-ECD is colored orange, hTIGIT-ECD is colored green.

Supplementary Movie 3: Representative MD simulation of the Fap2-ECD/TIGIT-ECD complex. 180 degree rotated view of SI Movie 2.

Supplementary Movie 4: Representative MD simulation of the Fap2-ECD/Gal-GalNAc-Thr complex. The movie shows a simulation over 50 nsec of the best docking pose from Autodock Vina. Gal-GalNAc-Thr is depicted gray, interacting residues of Fap2 are depicted violet.
